# Extensive ethnolinguistic diversity in Vietnam reflects multiple sources of genetic diversity

**DOI:** 10.1101/857367

**Authors:** Dang Liu, Nguyen Thuy Duong, Nguyen Dang Ton, Nguyen Van Phong, Brigitte Pakendorf, Nong Van Hai, Mark Stoneking

## Abstract

Vietnam features extensive ethnolinguistic diversity and occupies a key position in Mainland Southeast Asia (MSEA). Yet, the genetic diversity of Vietnam remains relatively unexplored, especially with genome-wide data, because previous studies have focused mainly on the majority Kinh group. Here we analyze newly-generated genome-wide SNP data for the Kinh and 21 additional ethnic groups in Vietnam, encompassing all five major language families in MSEA. In addition to analyzing the allele and haplotype sharing within the Vietnamese groups, we incorporate published data from both nearby modern populations and ancient samples for comparison. We find that the Vietnamese ethnolinguistic groups harbor multiple sources of genetic diversity that are associated with heterogeneous ancestry sharing profiles in each language family. However, linguistic diversity does not completely match genetic diversity; there have been extensive interactions between the Hmong-Mien and Tai-Kadai groups, and a likely case of cultural diffusion in which some Austro-Asiatic groups shifted to Austronesian languages. Overall, our results highlight the importance of genome-wide data from dense sampling of ethnolinguistic groups in providing new insights into the genetic diversity and history of an ethnolinguistically-diverse region, such as Vietnam.

## Introduction

Mainland Southeast Asia (MSEA) is of great interest in terms of ethnolinguistic diversity and deep population history. The early settlement of anatomically modern humans in MSEA dates back to at least 65 thousand years ago (kya) (Bae, et al. 2017; Demeter, et al. 2017), and is associated with the formation of a hunter-gatherer tradition called Hoabinhian (Higham 2013). Since the Neolithic period, which began about ∼4-5 kya, cultural transitions and diversification have happened multiple times (Edmondson and Gregerson 2007; Enfield, et al. 2011; Bellwood 2015, 2018; Habu, et al. 2018), eventually leading to the extraordinary cultural diversity in present day MSEA. To date, there are hundreds of ethnolinguistic groups in MSEA, speaking languages belonging to five major language families: Austro-Asiatic (AA), Austronesian (AN), Hmong-Mien (HM), Tai-Kadai (TK), and Sino-Tibetan (ST).

Vietnam occupies a key position in MSEA. It borders China, Laos, and Cambodia, and possesses a long coastline, allowing interactions with populations from southern China, MSEA, and Island Southeast Asia (ISEA). Vietnam has a population size of more than 96 million people (www.gso.gov.vn; accessed *the General Statistics Office of Vietnam* in September 2019), comprising 54 official ethnic groups; 110 languages are spoken in the country (Eberhard, et al. 2019). Most of these ethnic groups are found in either the southern highlands (mainly the AA and AN groups) or the northern highlands; the latter are especially heterogeneous and include AA, HM, TK, and ST groups (Eberhard, et al. 2019). The majority ethnic group in the lowlands is the AA-speaking Kinh, comprising ∼86% of the population (Dang, et al. 2016; Eberhard, et al. 2019), hence the genetic studies of Vietnamese to date have focused mainly on the Kinh (Vu-Trieu, et al. 1997; Ivanova, et al. 1999; Pischedda, et al. 2017; Le, et al. 2019). The genetic profiles of the other 53 official ethnic groups remain largely unexplored, leaving a substantial gap in our understanding of their genetic relationships and history.

The presence of five language families in Vietnam suggests diverse origins for this ethnolinguistic diversity. According to linguistic studies, the AA family is thought to be the oldest language family in MSEA, dating back ∼4-4.5 kya (Edmondson and Gregerson 2007; Enfield, et al. 2011; Bellwood 2015). Possible origins of the AA family include southern China, MSEA, or India (Bellwood 2015). The ST family originated in northern China (Sagart, et al. 2019; Zhang, et al. 2019), then moved southward into MSEA ∼3 kya (Bellwood 2015). The HM and TK families are thought to have originated in southern China, moving southward into MSEA in separate migrations ∼2.5 kya (Edmondson and Gregerson 2007; Bellwood 2015). The AN family expanded from Taiwan into ISEA, and from ISEA into MSEA, at about the same time (Bellwood 2015). The AN and TK proto-languages may be related, and early TK, HM, ST, and AA speakers may have encountered one another and interacted in what is now southern China, particularly the Southern Yunnan Interaction sphere (Enfield, et al. 2011; Habu, et al. 2018).

Cranial and facial skeleton evidence support a “two-layer” migration model for the settlement of MSEA, with the first arrival of hunter-gatherers during the Pleistocene followed by multiple waves of farmers from what is now southern China during the Neolithic period ∼4-5 kya (Matsumura and Oxenham 2014; Bellwood 2018). During the Bronze and Iron Age (∼1.5-2.5 kya), the Dong Son culture developed in northern Vietnam and spread across MSEA (Habu, et al. 2018). At about the same time, the Sa Huynh culture appeared in southern Vietnam and developed an extensive trade network that extended across MSEA (Bellwood 2015). The AN-speaking Cham people, who are thought to be associated with the Sa Huynh culture, arrived in south-central Vietnam, probably from Borneo, around 1.8-2.5 kya, and developed an extensive kingdom (Edmondson and Gregerson 2007; Bellwood 2015; Habu, et al. 2018). While the linguistic and archeological evidence thus suggests several population movements into MSEA including Vietnam, it is not clear to what extent these diverse language families spread by demic vs. cultural diffusion.

Genetic studies can inform on this question. For example, ancient genome studies have provided indications of demic diffusion, in that the present-day AA groups in MSEA show evidence of admixture involving Hoabinhian hunter-gatherers and the ancestors of Neolithic East Asians (Lipson, et al. 2018; McColl, et al. 2018). Another study of the mitochondrial DNA (mtDNA) control-region of the AN-speaking Cham demonstrated that they are likely to have resulted from language and culture shift of the indigenous AA-speaking Mon-Khmer populations to an AN language and culture (Peng, et al. 2010). A later study generated mtDNA control-region data from the Kinh and 4 ethnic minority groups and identified different haplogroup profiles among the AA, TK, HM, and ST groups (Pischedda, et al. 2017). More recent studies analyzed complete mtDNA genome sequences (Duong, et al. 2018; Macholdt, et al. 2019) and partial sequences of the male-specific portion of the Y chromosome (MSY) (Macholdt, et al. 2019) from the Kinh and 16 ethnic minority groups and further confirmed the diverse genetic profile in Vietnam. However, genome-wide studies, which can provide more resolution and additional insights into population relationships and history, are so far limited to the Kinh (Pischedda, et al. 2017; Le, et al. 2019). To further investigate the genetic diversity in Vietnam, we generated genome-wide SNP data from 22 Vietnamese ethnolinguistic groups, speaking languages that encompass all five families in MSEA. We incorporate published data, and analyze the allele and haplotype sharing within the Vietnamese groups and between them and both nearby modern populations and nearby SEA ancient samples. Our results provide new insights into the genetic diversity of these ethnolinguistically-diverse groups, including their recent interactions and demography.

## Results

### Overview of population structure

We genotyped 22 ethnolinguistic groups from Vietnam (fig. 1) and merged the data with data from nearby modern populations and ancient samples. We started by applying principle components analysis (PCA) and the clustering algorithm ADMIXTURE (Alexander, et al. 2009) to explore population structure. On a global scale, the strongest signal (i.e. variation along PC1) separates most Indian groups from the East Asian groups (fig. 2A, supplementary fig. S1 and table S1), with the Indian groups Kharia (#83 in the figures) and Onge (#82) placed between them. The ancient East Asian sample from Tianyuan (#1) and the Hoabinhian samples from Pha Faen (#2) and Gua Cha (#3) are projected between the Onge and the Jehai (#45) from Malaysia (fig. 2A, supplementary fig. S1 and table S1). The addition of PC2 further spreads out the East Asian groups, with northern Chinese groups (#67-73) at one end and ISEA groups (#45-58) at the other (fig. 2A, supplementary fig. S1 and table S1). With respect to language family, the ST, HM, and TK groups are mostly separated from AA and AN groups. Neolithic SEA ancient samples (#4-12) are mostly projected near the AA and AN groups, except that the sample from Oakaie (#7) is projected near the ST groups and other northern Chinese groups (fig. 2A, supplementary fig. S1 and table S1). The Bronze (#13) and Iron age (#14-15) samples are shifted more toward the present day Vietnamese (#19-40), especially the AA groups from Vietnam (#34-37), which are closer to the TK, HM, and ST groups than are the AA groups from other locations (fig. 2A, supplementary fig. S1 and table S1). The historical sample from Hon Hai Co Tien (#18) is projected near the Vietnamese AA and TK groups, while the samples from Supu Hujung (#16) and Kinabatagan (#17) are projected with the Taiwanese AN groups (#59-65) near the AN groups from ISEA and AA groups from Thailand (#42-43) (fig. 2A, supplementary fig. S1 and table S1).

**Fig. 1.**
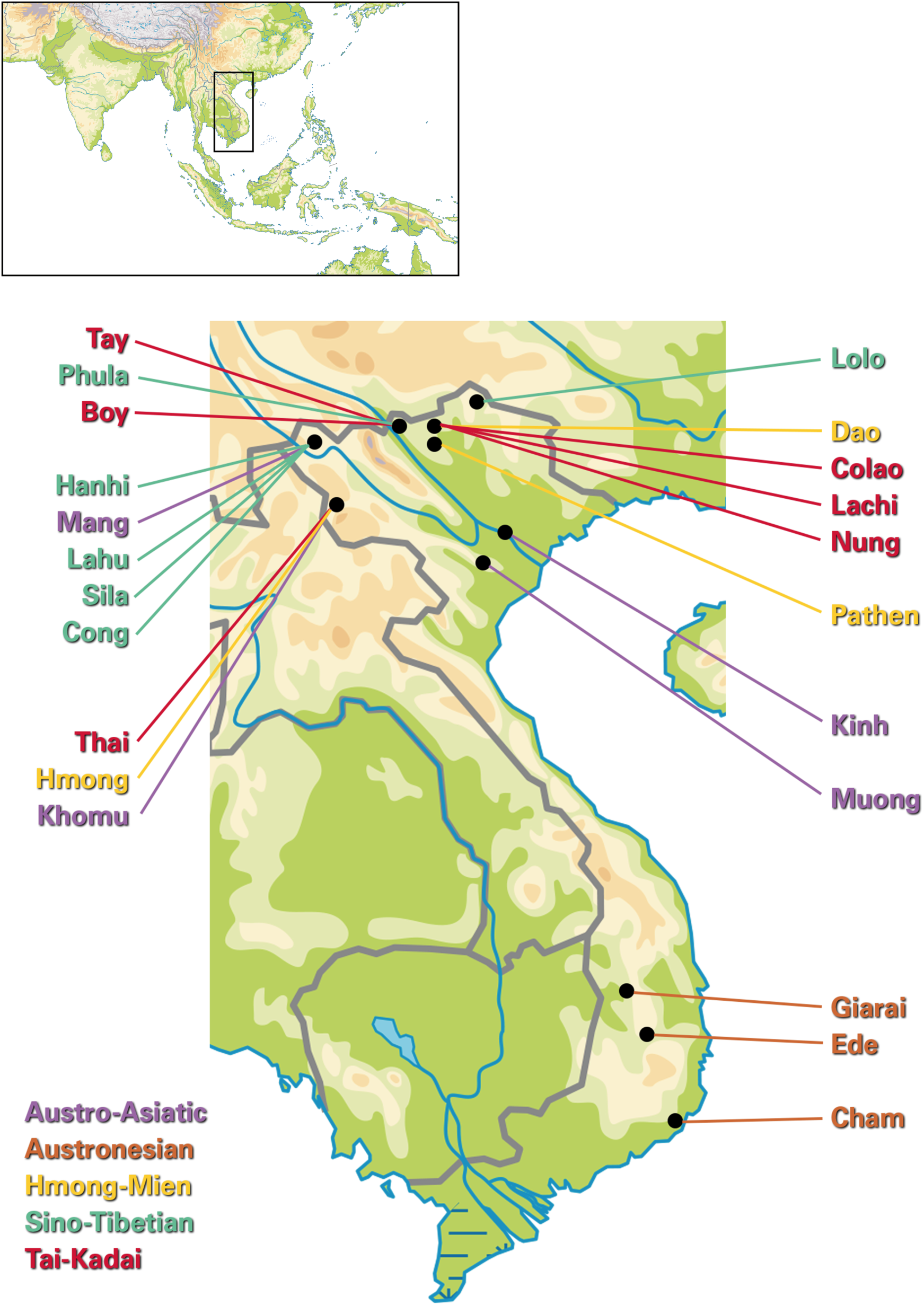
Map of the sampled Vietnamese ethnolinguistic groups. Dots denote the median of the sampling geographic coordinates per group.

**Fig. 2.**
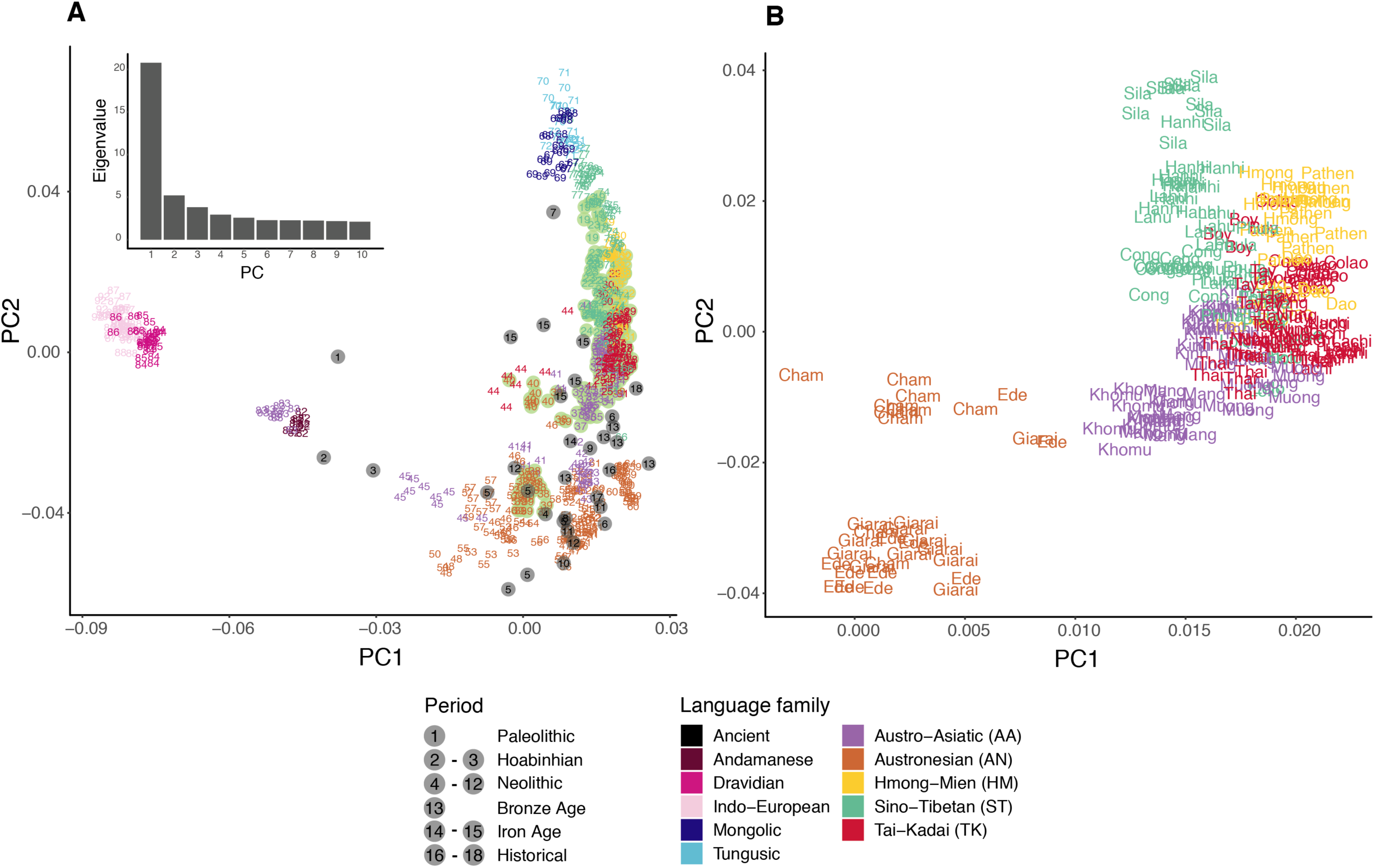
PCA analyses. (A) PCA analysis of 712 individuals and 33,666 SNPs, with individuals colored according to language families. The eigenvalues from PC1 to PC10 are shown in the top left corner. Ancient samples are shown as grey dots, while the present-day Vietnamese are shown as light green dots. Each population and ancient sample is numbered according to supplementary table S1. (B) Vietnamese populations only, zoomed-in from (A).

Within modern Vietnamese groups, individuals from the same language family are mostly placed together (fig. 2B). There is some overlapping of individuals from different language families, except that AN groups are distinct from the others, closer to the AA-speaking Cambodian (#41), Htin Mal (#42), Mlabri (#43), and many ISEA populations, all of whom speak AN languages (fig. 2A and supplementary table S1). Within the AN groups, the Cham are mostly separated from the Ede and Giarai along PC2 (fig. 2B). When considering additional PCs, the Ede and Giarai are strongly differentiated on PC3 from all other groups, except for the AA-speaking Khomu and Mang (supplementary fig. S2). Additional PCs tend to highlight the distinctiveness of the Mang, ST-speaking Sila, and TK-speaking Colao and Lachi (supplementary fig. S2).

We next performed an ADMIXTURE analysis and found that the lowest cross validation error occurs at K = 6 (supplementary fig. S3). Under the model of K = 6, there is: a brown source present only in the Mbuti; a pink source enriched in both the French and Indian groups; a blue source enriched in AA-speaking groups and in AN-speaking groups from Indonesia, Malaysia and Vietnam; a black source enriched in AN-speaking groups from Taiwan, Philippines, and Indonesia; a purple source appearing in all of the Chinese groups and enriched in the Vietnamese ST groups; and a dark green source absent before K = 6 appearing in the southern Chinese groups and enriched in Vietnamese HM and TK groups (fig. 3 and supplementary fig. S4). In general, Vietnamese groups show diverse genetic profiles with variable amounts of the dark green, purple, blue, and black sources (fig. 3). The Vietnamese AN groups are notable in that the amount of Austronesian-related black source in them does not surpass other Vietnamese groups, and in having higher frequencies of the pink source (12% on average in AN groups compared to a maximum of 1.5% in the other Vietnamese groups; supplementary table S2). The Ede and Giarai show less purple source than the Cham (3% vs. 13%; supplementary table S2).

**Fig. 3.**
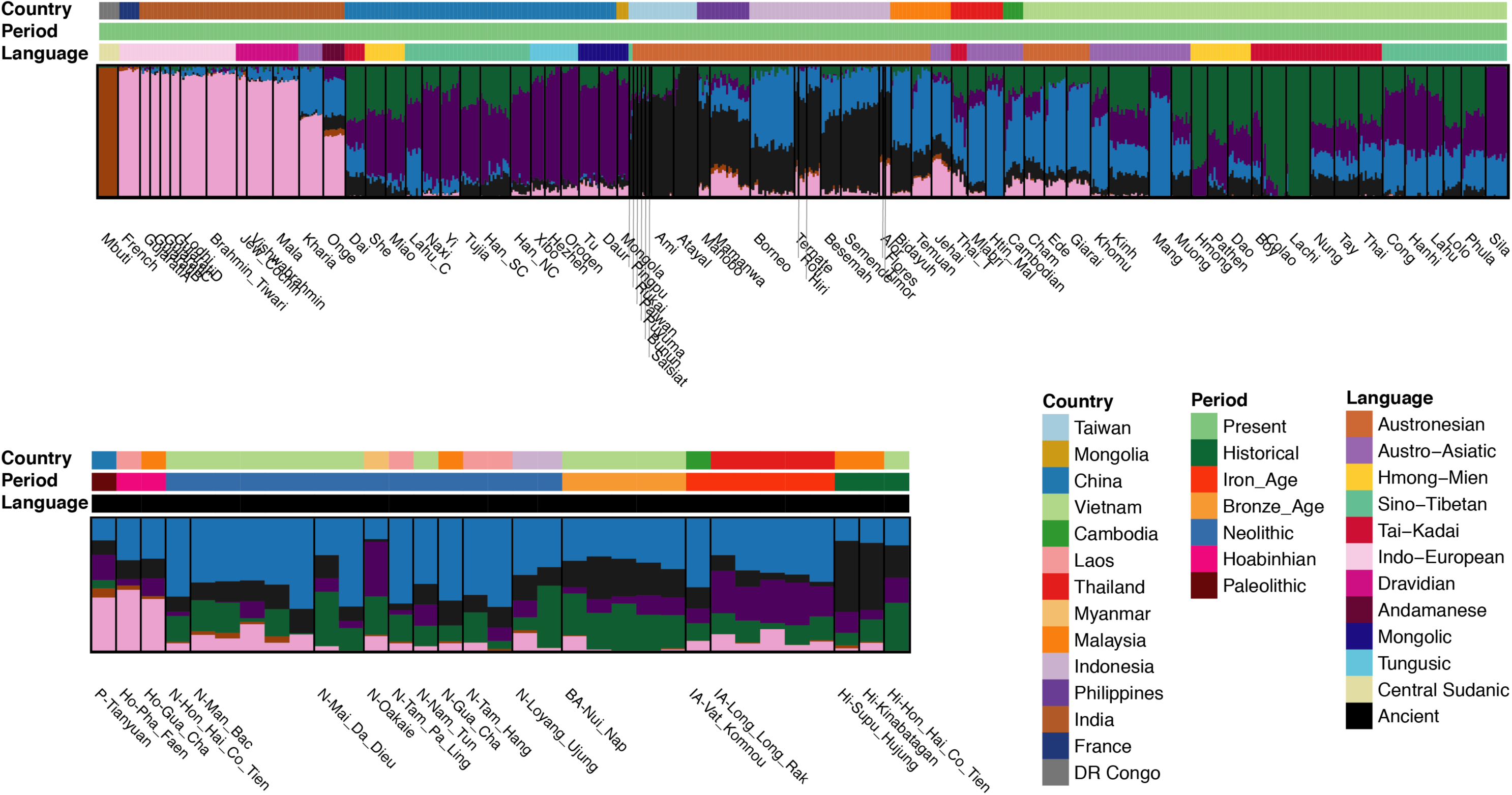
ADMIXTURE analyses. ADMIXTURE result for K = 6, which minimized the cross-validation error (supplementary fig. S3). Regions are labeled at the top, while population/ancient sample names are labeled at the bottom. The color bar denotes the language families and time periods for the modern and ancient samples, respectively.

With respect to ancient samples, the pink source is enriched in the Tianyuan and Hoabinhian samples, while the blue source is enriched in the Neolithic samples (fig. 3 and supplementary table S2). The blue source decreases in ancient samples younger than the Neolithic, with a concomitant increase of the green, purple, or black sources (fig. 3). Specifically, the green and black sources increase in the Bronze Age and historical samples in Vietnam (fig. 3 and supplementary table S2). The black source also increases in the historical samples from Malaysia, while the purple source is enriched in the Iron Age sample from Long Long Rak (Thailand) but also in the Neolithic sample from Oakaie (Myanmar; fig. 3 and supplementary table S2).

Overall, the Vietnamese AN groups are more similar in this analysis to some groups from Malaysia, Thailand, and Cambodia, and to the Neolithic ancient samples (except for the sample from Oakaie), than to other Vietnamese groups (fig. 3). The HM-speaking Hmong and the TK-speaking Colao and Lachi stand out in lacking the black and blue sources, and in general both HM and TK groups present the highest frequencies of the green source. The black source is relatively low in ST groups, and the Sila differ from other ST groups in lacking the black and green sources; in this respect the Sila are more similar to the AA-speaking Mang (but with more of the purple source and less of the blue source than in the Mang) (fig. 3). The remaining Vietnamese groups present fairly similar profiles (albeit with some variation in the frequencies of specific sources) that are also similar to the Dai from southern China (fig. 3).

Although higher values of K are associated with higher cross-validation errors, they can nevertheless provide additional insights. At K = 7, the French get their own source, which is practically absent in all of the Vietnamese individuals (supplementary fig. S4), and confirms that the pink source in the Vietnamese AN-speaking groups is likely shared ancestry with Indian groups. At K = 8, many ISEA populations have high frequencies of the peach source, which is at highest frequency in Alor and Timor (supplementary fig. S4). This source decreases the black source present in the SEA groups, except for the Vietnamese AA, TK, HM, and ST groups. At K = 9 and 10, the Mang and Lachi get their own source, respectively (supplementary fig. S4). At K = 11, the Sila obtain their own source, which also shows up in the ST groups in both China and Vietnam (supplementary fig. S4). At K = 12 and 13, the Atayal and Colao get their own source, respectively (supplementary fig. S5). At K = 14, the Htin Mal get their own source, which also shows up in many Neolithic samples (supplementary figs. S4 and S5). Finally, at K = 15, the Hmong get their own source, which is also present in all of the HM groups (supplementary fig. S4). Therefore, out of 9 additional K-values (K = 7-15), 5 involve Vietnamese populations getting their own source, which is a further indication of the extensive genetic structure among Vietnamese groups.

### Investigation of population relationships and demography

The above analyses (PC and ADMIXTURE) are descriptive analyses that provide an overview of the relationships of the populations analyzed. To further explore and quantify these relationships, we used outgroup *f3* and *f4* statistics to identify ancestry sharing based on allele sharing, and identity by descent (IBD) approaches to investigate demography and recent contact based on haplotype sharing.

#### Outgroup f3

Higher values of the outgroup *f3* statistic indicate more shared drift, and hence a closer relationship, between two test populations since their divergence from the outgroup population. We first compared the *f3* results within Vietnamese groups. The AN groups are again most distant from others (fig. 4 and supplementary fig. S6), and also show more shared drift with some non-AN groups than with each other. The AA groups exhibit two distinct sharing profiles: the Mang/Khomu have relatively low levels of shared drift with all other Vietnamese groups, while the Muong/Kinh have higher levels of sharing with each other and with some TK, HM, and ST groups (fig. 4 and supplementary fig. S6). The TK and HM groups share the most with each other, with Muong/Kinh, and with the ST-speaking Lolo and Phula (fig. 4 and supplementary fig. S6). Next, we investigated the relationships between Vietnamese and neighboring modern populations. Vietnam ethnolinguistic groups overall tend to show the closest relationships with Taiwanese and southern Chinese groups (fig. 4 and supplementary fig. S7). Most of them show top *f3* values with the AN-speaking Ami and TK-speaking Dai (fig. 4 and supplementary fig. S7). Consistent with the PCA results (supplementary fig. S1), the Vietnamese groups are mostly distant from Indian populations (fig. 4 and supplementary fig. S7). The AN groups, especially Ede and Giarai, exhibit higher *f3* values with the AA-speaking Mlabri and Htin Mal, while Cham shows more sharing with the AN-speaking Ami and TK-speaking Dai (fig. 4 and supplementary fig. S7). The AA groups can again be separated into the Mang/Khomu vs, Muong/Kinh, with the former overall showing lower levels of sharing, but relatively more sharing with the AA-speaking Htin Mal and Mlabri, than the latter; the Muong/Kinh share relatively more with the AN-speaking Ami and TK-speaking Dai (fig. 4 and supplementary fig. S7). For the HM groups, the Hmong and Pathen show higher *f3* values with the HM-speaking Miao and She, and the Pathen as well as Dao also show higher *f3* values with the AN-speaking Ami and TK-speaking Dai (fig. 4 and supplementary fig. S7). All of the TK groups display higher *f3* values with the Ami, while the Boy, Lachi, and Thai share the most with the TK-speaking Dai (fig. 4 and supplementary fig. S7). The HM and TK groups generally seem to share more with the TK-speaking Dai, HM-speaking Miao and She, ST-speaking Tujia and Han, and the AN-sepaking Ami and Atayal, than with the Vietnamese AA, AN, and ST groups (fig. 4 and supplementary fig. S7). The ST groups exhibit high *f3* values with several southern Chinese populations, and the Hanhi, Lahu, and Phula show especially higher sharing with the ST-speaking Chinese Lahu (fig. 4 and supplementary fig. S7). Similar outgroup *f3* profiles are obtained when the French are used as an outgroup instead of the Mbuti (supplementary fig. S8).

**Fig. 4.**
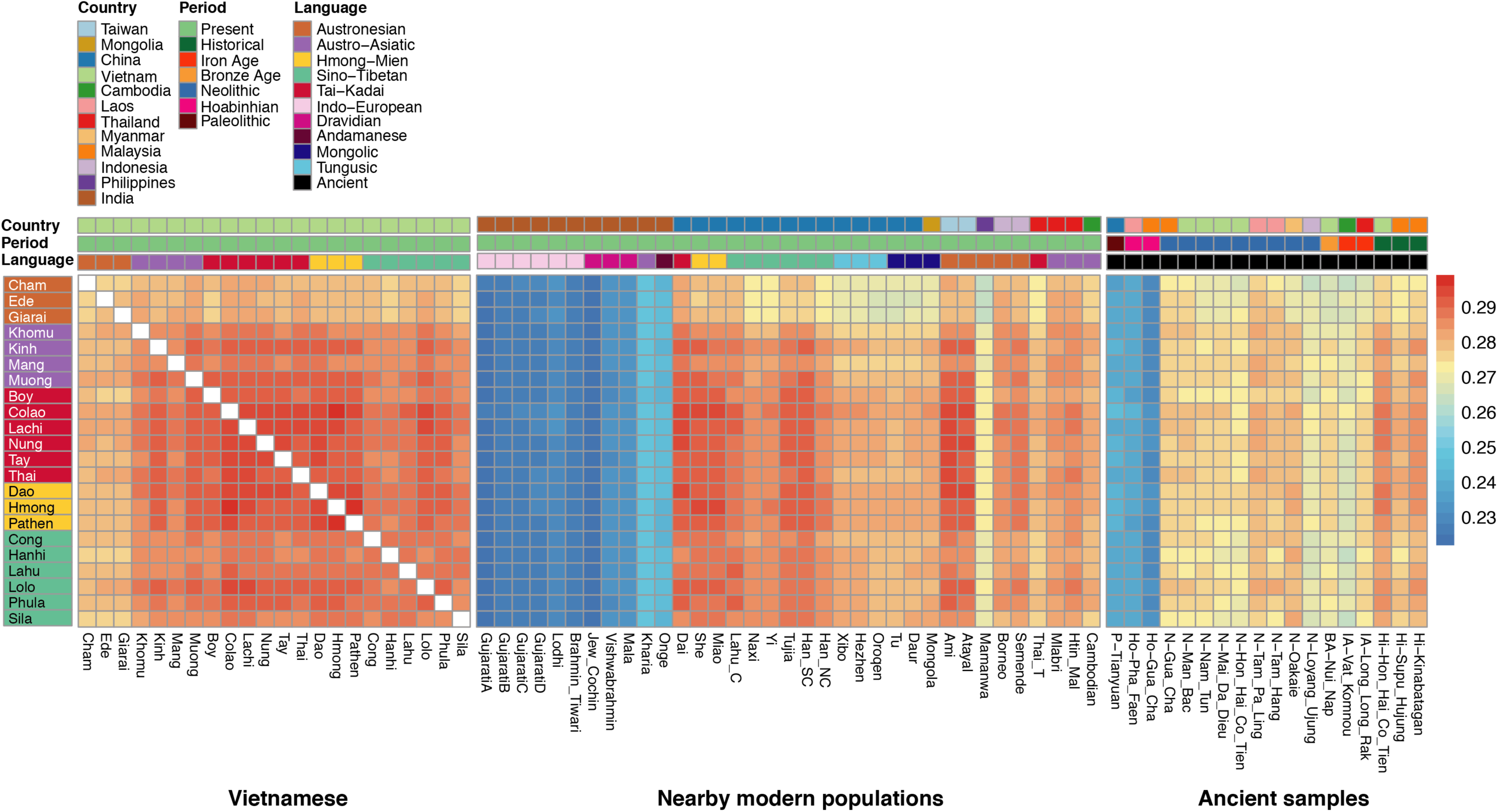
Heatmap of outgroup *f3* profiles. A heatmap based on the values of the *f3* statistic for all pairs of populations/ancient samples, using Mbuti as the outgroup. Shown are: comparisons among the Vietnamese groups (left); with the nearby modern populations (middle); and with the ancient samples (right). The three different color bars at the top denote separately the countries, time periods, and language families, according to the key. The Vietnamese group labels are also shaded according to language family.

When compared to ancient samples, all the Vietnamese groups exhibit high *f3* values with the historical samples from Hon Hai Co Tien (Vietnam) and Kinabatagan (Malaysia), except for the AN groups (fig. 4). The normalized *f3* values tend to be especially high (> 0.95) with the historical sample from Hon Hai Co Tien (fig. 4 and supplementary fig. S9). The AN-speaking Ede/Giarai as well as the AA-speaking Mang/Khomu show higher *f3* values with the Neolithic samples from Tam Pa Ling (Laos), Tam Hang (Laos), Gua Cha (Malaysia), and Man Bac (Vietnam; fig. 4 and supplementary fig. S9). The *f3* values with the Neolithic sample from Oakaie (Myanmar), Bronze Age sample from Nui Nap (Vietnam), and Iron Age sample from Long Long Rak (Thailand) are generally high with all groups except the AN groups (fig. 4 and supplementary fig. S9). The smallest *f3* values are those with the Paleolithic sample from Tianyuan and the Hoabinhian samples, and the *f3* values with these samples show little variation among Vietnamese groups (fig. 4 and supplementary fig. S9).

#### IBD

We next investigated interactions within/between populations within the past ∼3 kya by analyzing IBD (Ralph and Coop 2013; Al-Asadi, et al. 2019). The number and length of IBD segments shared within a population provides further insights into population demography (Browning and Browning 2015; Browning, et al. 2018; Ceballos, et al. 2018; Severson, et al. 2019). The Hmong, Pathen, Lachi, Boy, Colao, Mang, Lolo, and Sila all show elevated levels of within population IBD sharing, while the Kinh have the lowest level (supplementary fig. S10). We used the IBD sharing within each population to directly estimate recent changes in effective population size, i.e. within the past 50 generations (Browning and Browning 2015). The Boy, Lachi, Dao, Sila, Cong, and Khomu are inferred to have experienced bottleneck events, while the AA-speaking Kinh and Muong have undergone population expansions beginning around 15-20 generations (∼450-600 years) ago (fig. 5). The three AN groups have also undergone a slight reduction in population size ∼450-600 years ago, followed by population expansions ∼300-450 years ago (fig. 5). Other populations show no obvious bottleneck events but an overall decrease in size; in particular, the Colao, Hmong, Lolo, and Mang have very small effective population sizes (fig. 5).

**Fig. 5.**
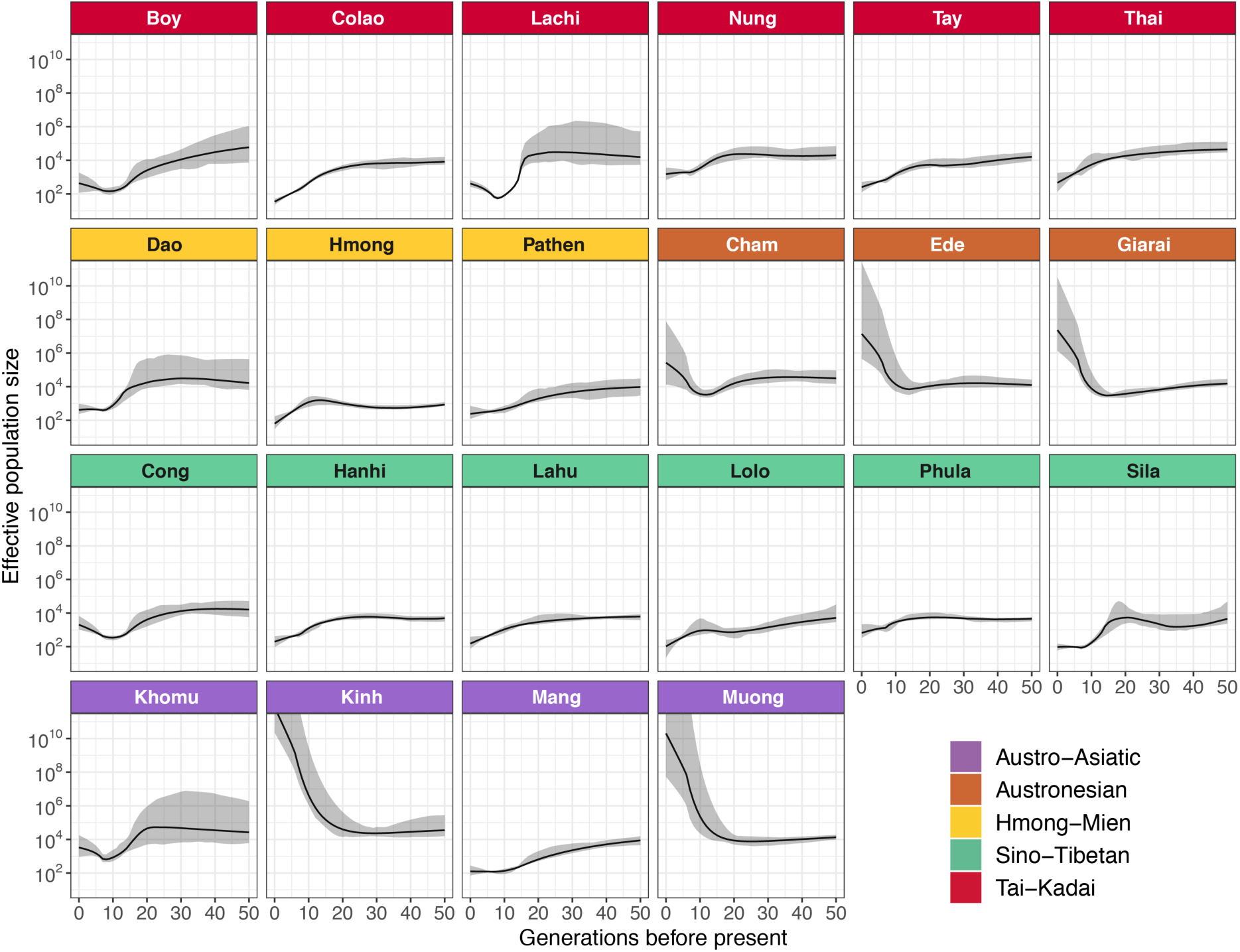
Effective population size of Vietnamese ethnolinguistic groups over the past 50 generations. Each panel depicts an ethnolinguistic group. The panels are colored according to language family; 95% confidence intervals are shaded in grey.

While IBD sharing within populations provides insights into population size changes, IBD sharing between populations provides insights into recent contact. More shared IBD blocks between populations implies more interaction, and the longer the shared IBD blocks, the more recent the interaction. As the IBD blocks will be broken down by chromosomal recombination through time, we can thus infer the time of interaction(s) based on the range of block lengths. We analyzed IBD blocks in three categories: 1-5 cM, corresponding to 90 generations ago, 5-10 cM, corresponding to 23 generations ago; and >10 cM, corresponding to 7.5 generations ago (Al-Asadi, et al. 2019). The IBD results show wider interaction of Vietnam ethnolinguistic groups with neighboring populations and within their language families 90 generations ago (∼2.7 kya), becoming more and more localized by 23 (∼0.7 kya) and 7.5 (∼0.2 kya) generations ago (fig. 6). In the range of 1-5 cM, both the Vietnamese and neighboring populations within the same language families tend to be linked together, except that the Vietnamese AN groups share IBD blocks with AA-speaking groups (fig. 6). All of the Vietnamese ST groups have links to the ST-speaking Chinese Lahu, while the Phula and Lolo have more connections with the HM and TK groups (fig. 6). The HM groups interact with the HM-speaking Miao, She, and the TK groups (fig. 6). The strongest between-group sharing is between Hmong and Colao (fig. 6). The Hmong also show interactions with the ST groups (fig. 6). The Thai and Nung from the TK groups interact with the TK-speaking Dai (fig. 6). The Khomu from the AA groups interact with the AN groups (fig. 6). The AN groups together with the AA groups have interactions with the AA speaking Mlabri, Htin Mal, Cambodian, which further connect to Borneo and other AN-speaking populations (fig. 6) In the range of 5-10 cM, the only sharing between Vietnamese and others is Vietnamese Lahu with Chinese Lahu, and Hmong with Miao (fig. 6). The HM and ST groups still have strong sharing with the HM-speaking Miao and ST-speaking Chinese Lahu, respectively (fig. 6). While the HM, TK, and ST groups are intermixed, the AN groups share exclusively with each other (fig. 6). Sila, Hanhi, and Lahu also exhibit sharing only within the ST groups (fig. 6). In the range of over 10 cM, sharing is limited to only a few localized pairs between ST, HM, and TK groups as well as within the AN groups (fig. 6). The ST-speaking Sila and Hanhi exhibit strong IBD sharing with each other over the entire size range of IBD blocks (fig. 6). Notably, the AA-speaking Kinh and Muong do not share any IBD blocks with any other group, irrespective of the size of the blocks.

**Fig. 6.**
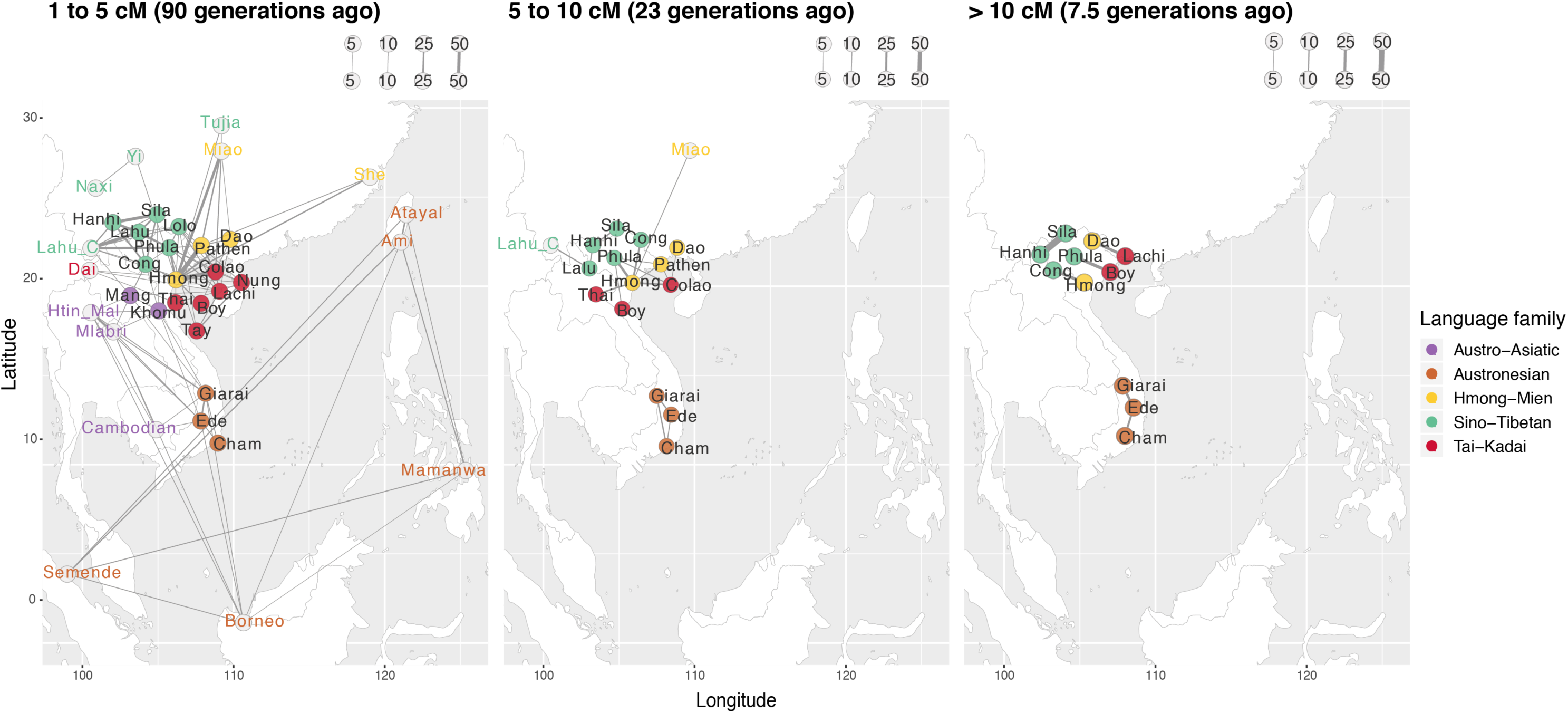
IBD sharing between populations. Network visualizations of the mean of summed IBD lengths shared between populations, with identified IBD blocks in the range of 1 to 5 cM (90 generations ago/ ∼2.7 kya), 5 to 10 cM (23 generations ago/ ∼0.7 kya), and over 10 cM (7.5 generations ago/ ∼0.2 kya). We only show the sharing involving Vietnamese groups. The signals were enriched by requiring an average of at least 2 shared IBD blocks per pair of individuals (4 for the range of 1 to 5 cM) as illustrated by the pink rectangles (chromosome) with purple blocks (IBD blocks). Each node stands for a population, and each edge indicates the IBD sharing between populations. The nodes of the Vietnamese groups are colored according to language family. The width of each edge is proportional to the mean of the summed IBD length, with the scale (cM) provided in the top-right portion of each figure. Also provided is a map of the populations in the IBD network, which is also colored by language family.

#### *f4* statistics

We further investigated the relationships of Vietnamese groups with representative source populations for each language family. Based on the f3 and IBD sharing results, we selected the Htin Mal (AA), Atayal (AN), Miao (HM), Dai (TK), and Chinese Lahu (ST) as the representative source populations for the five language families in Vietnam. We then calculated f4 statistics of the form *f4*(Source populations, southern Han Chinese; Vietnamese, Mbuti) to test if each Vietnamese group shares any excess ancestry with any of the representative source populations, compared to the southern Han Chinese. Significantly positive Z-scores indicate excess shared ancestry between the Vietnamese group and the source population, while significantly negative Z-scores indicate excess shared ancestry between the Vietnamese group and southern Han Chinese. The resulting Vietnamese *f4* profiles are heterogeneous within each language family (fig. 7). The AA-speaking Khomu and AN-speaking Ede and Giarai show significant excess ancestry sharing with the AA-speaking Htin Mal. All other Vietnamese groups show excess ancestry sharing with southern Han Chinese, except for the AA-speaking Mang and the AN-speaking Cham, which show no excess shared ancestry (fig. 7). With the AN-speaking Atayal as the source, the only significant sharing is between Atayal and the AN-speaking Ede and Giarai, and between southern Han Chinese and the ST-speaking Sila (fig. 7). With the HM-speaking Miao as the source, the only significant sharing is between the HM-speaking Hmong and the Miao (fig. 7). With the TK-speaking Dai as the source, there is significant sharing between the Dai and the ST-speaking Lolo, the TK-speaking Thai and Lachi, the AA groups except the Kinh, and all of the AN groups (fig. 7). Finally, with the ST-speaking Chinese Lahu as the source, there is significant sharing between them and the ST-speaking Vietnamese Lahu. In contrast, the southern Han Chinese share ancestry with all of the HM groups, all of the TK groups (except Lachi), and with the AA-speaking Muong and Kinh. (fig. 7).

**Fig. 7.**
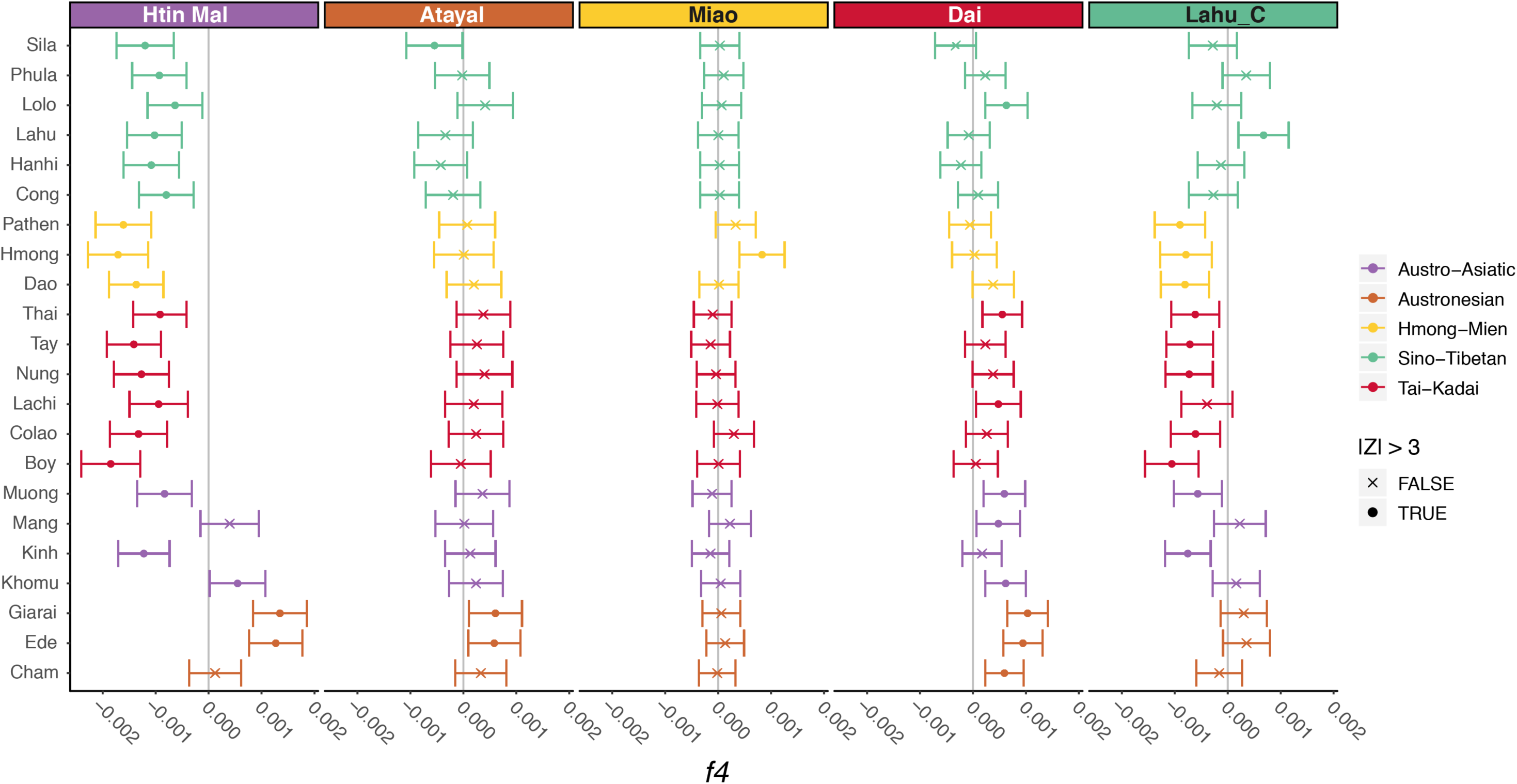
*f4* statistics comparing Vietnamese groups to representative source populations. Z-scores are for *f4*(W, southern Han Chinese; Y, Mbuti), where W is the source population (panel labels) and Y is the Vietnamese group (label on the Y axis). The vertical grey lines denote 0. The panels are colored according to language family.

When we used ancient samples as the source population in this *f4* statistic, no Vietnamese group shares excess ancestry with any ancient sample (supplementary fig. S11). Instead, practically all of the Vietnamese groups share excess ancestry with southern Han Chinese (supplementary fig. S11); the few exceptions, in which there is no excess sharing between the Vietnamese group and either the southern Han Chinese or the source population, involve various of the AN groups, Khomu, Mang, and/or Sila with the Neolithic samples and the Iron Age sample from Vat Komnou. Also, many Vietnamese groups share no excess ancestry with southern Han Chinese in the comparisons with historical samples (supplementary fig. S11).

The population structure analyses suggested a shift in the affinities of the ancient samples, with pre-Neolithic/Neolithic samples more similar to AA and AN groups, and more recent samples exhibiting more similarities to TK, HM, and ST groups (figs. 2 and 3). To further investigate this, we used Mlabri, Htin Mal, Borneo, Ami, and Mamanwa as a combined representative source of the AA and AN groups, and Dai, Miao, Chinese Lahu, southern Han Chinese, and northern Han Chinese as a combined representative source of the TK, HM, and ST groups, and then computed f4 statistics of the form *f4*(TK/HM/ST, AA/AN groups; Ancient samples, Mbuti). We found that the AA and AN groups indeed shared excess ancestry with the Hoabinhian sample from Pha Faen, most of the Neolithic samples except for the samples from Oakaie (which shares excess ancestry with the TK, HM, and ST groups) and Nam Tun, and the historical samples from Supu Hujung and Kinabatagan (supplementary fig. S12). This result supports the shift in affinities of ancient samples that was observed in the population structure analyses (figs. 2 and 3). Restricting the analyses with the ancient samples to transversions reduces the number of SNPs from 361,327 to 64,126, and correspondingly many of the Z-scores become non-significant (supplementary figs. S13 and S14).

### Admixture graph inference

Based on the sharing profiles revealed by the *f3,* IBD, and *f4* analyses, we next built admixture graphs for Vietnamese groups from each language family. Admixture graphs, which depict a history of population divergence and admixture events, use either a combination of *F*-statistics or a covariance matrix of the allele frequencies (Nielsen 2018). We first applied TreeMix (Pickrell and Pritchard 2012) and AdmixtureBayes (Nielsen 2018) to survey the potential admixture graphs based on the covariance matrix of allele frequencies, and then we further tested if these graphs are accepted in qpGraph (Patterson, et al. 2012), using a combination of *F*-statistics. Before building the graph for each language family, we first built a global tree with all the Vietnamese groups, the representative source populations used in the *f4* analyses, the Onge, selected ancient samples, and the Mbuti as an outgroup. We found that all of the ancient samples fall outside the Vietnamese clade, except that the historical sample from Kinabatagan shares an ancestor with the clade of the ST groups and an admixture source from the lineage leading to the AN-speaking Atayal (supplementary fig. S15). The AN groups are placed outside the clade of other Vietnamese groups; the former is close to the Neolithic samples from Tam Pa Ling and the AA-speaking Htin Mal (supplementary fig. S15). The AA-speaking Kinh and Muong and the ST-speaking Phula and Lolo are close to the HM and TK groups rather than to other groups from the same language family (supplementary fig. S15). The HM-speaking Dao is closer to the TK groups compared to other HM groups, while the TK-speaking Colao is placed in the clade of HM groups (supplementary fig. S15).

On a local scale, we started with a backbone graph with the representative source populations used in the *f4* analyses, the Onge, and the Mbuti as an outgroup, for further investigating the admixture graphs by each language family. The backbone graph shows that the first split separates the Onge from a branch leading the ST-speaking Chinese Lahu and the HM-speaking Miao (fig. 8A; TreeMix results in supplementary fig. S16). All other groups are derived via admixture events. The AA-speaking Htin Mal has ∼4% ancestry from the ancestor of the Onge and 96% ancestry from an ancestor of the Chinese Lahu and Miao (fig. 8A). The AN-speaking Atayal have ∼1% ancestry from this same Onge ancestor, and 99% ancestry from a source related to the Miao. Finally, the TK-speaking Dai have ∼85% ancestry from this same Miao-related source, and ∼15% ancestry from an ancestor of the Htin Mal (and thereby also share some ancestry with Onge and Atayal).

**Fig. 8.**
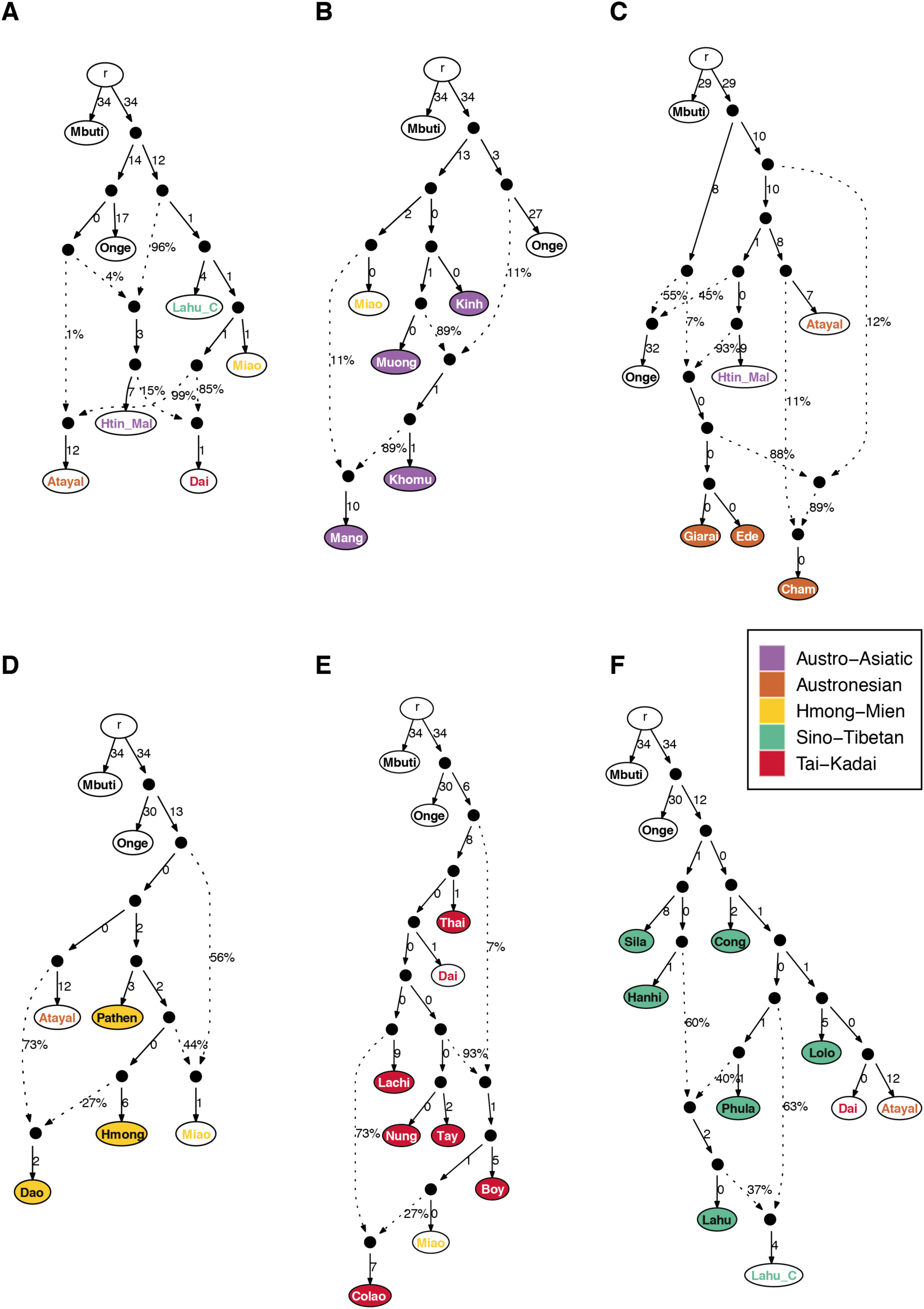
Admixture graphs of the Vietnamese groups, for each language family. The best-fitting admixture graphs are shown for the backbone populations and for the Vietnamese groups, done separately for each language family. The node r denotes the root. White nodes denote backbone populations. Backbone population labels and Vietnamese nodes are colored according to language family. Dashed arrows represent admixture edges, while solid arrows are drift edges reported in units of FST×1,000. (A) backbone populations (worst-fitting Z = −2.263). (B) AA groups (worst-fitting Z = −4.487; worst-fitting Z = 1.509 when removing the Kinh). (C) AN groups (worst-fitting Z = −1.258). (D) HM groups (worst-fitting Z = −1.462). (E) TK groups (worst-fitting Z = −3.151; worst-fitting Z = 2.721 when removing the Colao). (F) ST groups (worst-fitting Z = 3.499; worst-fitting Z = −2.106 when removing the Cong).

The admixture graph for the Vietnamese AA groups (fig. 8B; Treemix results in supplementary fig. S17) supports the division noted in previous analyses for the Kinh/Muong vs. the Khomu/Mang. The former share an ancestor with the Miao, while the latter are admixed from sources related to the Onge and the Muong (similar to the Htin Mal in the backbone graph), with the Mang in addition having ∼11% Miao-related ancestry. The AN groups show different histories for the Giarai and Ede vs. the Cham (fig. 8C; TreeMix results in supplementary fig. S18). The Giarai/Ede have ∼7% ancestry from an ancestor of the Onge, and ∼93% ancestry from an ancestor of the Htin Mal, while the Cham have ancestry from an ancestor of the Atayal and Htin Mal, an ancestor specifically of the Atayal, and an ancestor of the Giarai/Ede (thereby contributing Onge-related and additional Htin Mal-related ancestry). For the HM groups, the Hmong and Pathen share an ancestor with the Atayal, while the Dao is admixed from an ancestor of the Atayal and an ancestor of the Hmong (fig. 8D; TreeMix results in supplementary fig. S19). In this graph the Miao are modeled as having admixed ancestry from an ancestor of the Atayal/Pathen/Hmong and an ancestor of the Hmong. The TK groups Thai, Lachi, Nung, and Tay form a clade with the Dai (fig. 8E; TreeMix results in supplementary fig. S20). The Boy have admixed ancestry with an ancestor of this clade and an ancestor of the Nung and Tay, while the Colao have admixed ancestry involving an ancestor of the Miao (who in turn are closely related to the Boy) and an ancestor of the Lachi. All of the ST groups (except the Lahu) form a clade, together with the Dai and Atayal, with the Lolo most closely related to the Dai and Atayal (fig. 8F; TreeMix results in supplementary fig. S21). The Vietnamese Lahu have admixed ancestry from an ancestor of the Hanhi and an ancestor of the Phula (fig. 8F); in this analysis, the Chinese Lahu are modeled as having admixed ancestry from an ancestor of the Vietnamese Lahu and the Phula (fig. 8F).

## Discussion

### Extensive genetic diversity among Vietnamese groups

In this study, we have generated and analyzed genome-wide SNP data from 22 ethnolinguistic groups in Vietnam encompassing all five language families in MSEA (supplementary table S1). We found extensive genetic diversity among Vietnamese groups in the PCA and ADMIXTURE analyses, especially in additional components where many isolated groups stand out (figs. 2, 3, supplementary figs. S2 and S4). Hence, the majority group Kinh, which have been the focus of previous studies, may not reflect the total Vietnamese diversity, although we note that our sample of Kinh is relatively small and may not reflect the true genetic diversity of the Kinh.

Overall, the AN groups are distinct from the others but closest to the AA groups (fig. 2). The HM, TK, and ST groups share more ancestry with present-day southern Chinese groups, and the former two are more closely related to each other (figs. 2-4 and 6). By incorporating ancient samples from SEA and China, we have shown that the AA ancestry rose in the Neolithic period, followed by an increase of AN, HM/TK, or ST ancestry (according to the region) in later periods (fig. 3 and supplementary table S2). This population turnover from the Neolithic to later periods, with additional Chinese-related ancestry, is consistent with the archeological and linguistic studies (Edmondson and Gregerson 2007; Bellwood 2015; Habu, et al. 2018), but contradicts a previous study, based on much more limited sampling, that claimed a largely indigenous origin for Vietnamese groups (Le, et al. 2019). As a result, the overall Vietnamese genetic diversity likely reflects multiple waves of ancestry since the Neolithic period, which correlate somewhat (but not completely) with the language families, as we now discuss for each language family.

### Austro-Asiatic

The possible origins of the AA family include southern China, MSEA, or India (Bellwood 2015). It is thought to be the oldest language family in MSEA, which emerged after the Hoabinhian tradition ∼4-5 kya (Bellwood 2015). Ancient genome studies have suggested that the present-day AA groups in MSEA are descendants of Hoabinhian hunter-gatherers and ancestral East Asians from southern China admixing during the Neolithic farming expansion (Lipson, et al. 2018; McColl, et al. 2018). Consistent with this scenario, we find that the indigenous AA groups Htin Mal and Khomu have 4% and 11% ancestry from the Hoabinhian hunter-gatherers and 96% and 89% ancestry from the ancestral East Asians, respectively (figs. 8A and 8B). The AA-speaking Mang are closer to the Khomu compared to the Kinh and Muong, but they also share ancestry with the ST-speaking Lahu in the TreeMix analysis (supplementary fig. S17), and they share ancestry with the HM-speaking Miao in the qpGraph analysis (fig. 8B). This ancestry sharing with ST-speaking Chinese Lahu could reflect the close proximity of the Mang to ST groups (fig. 1). In contrast, the AA-speaking Kinh and Muong share more drift with HM and TK groups than with other AA-speaking groups (fig. 4). In particular, they are not estimated as having ancestry from the Hoabinhians in the admixture graph, in contrast to the Mang and Khomu (fig. 8B and supplementary fig. S17). This is consistent with previous suggestions that the Kinh and Muong may be related to the Dong Son culture and have ancestors from southern China (Dang, et al. 2016; Habu, et al. 2018), but contradicts one recent study stating that the Kinh appear to be an indigenous SEA group with less EA ancestry (Le, et al. 2019). However, this study included only the Kinh and Thai for the SEA groups and the Han, Korean, and Japanese for the EA groups. It is likely that our inclusion of many more SEA and Chinese groups, and more detailed sampling of Vietnamese ethnolinguistic groups, provides a more accurate picture of their relationships.

As the Kinh and Muong have the highest census size of Vietnamese groups (Dang, et al. 2016; Eberhard, et al. 2019), it seems likely that they have interacted extensively with each other as well as with HM and TK groups. However, while we found that the Khomu and the Mang share IBD blocks with each other and with ST and AN groups, we did not find any strong IBD sharing between the Kinh and Muong and other groups (fig. 6). Moreover, we observed exponential population expansions in the Kinh and Muong, compared to population contractions in the Khomu and Mang, ∼20 generations (∼600 years) ago (fig. 5). This expansion may have accelerated the breaking down of shared IBD blocks by increasing recombination events, and hence the Kinh and Muong may have had some recent contact with HM and TK groups, even if this is not visible in the IBD-sharing analysis.

### Austronesian

The origin of the AN family is proposed to be Taiwan (Gray, et al. 2009; Ko, et al. 2014; Bellwood 2015). The expansion of the AN groups into ISEA is dated ∼3-4 kya (Gray, et al. 2009; Bellwood 2015), while the emergence of the AN family in MSEA is thought to have happened ∼2.5 kya (Peng, et al. 2010; Bellwood 2015). Previous linguistic studies thus suggested that the introduction of the AN family into MSEA was via migration from ISEA after the initial expansion from Taiwan (Edmondson and Gregerson 2007; Enfield, et al. 2011; Bellwood 2015). In particular, the ancestors of the Cham are thought to have come from ISEA, probably Indonesia, and they established the Kingdom of Champa and dominated southern Vietnam during the 2^nd^ to mid-15^th^ century (Edmondson and Gregerson 2007; Enfield, et al. 2011; Bellwood 2015; Habu, et al. 2018). In contrast, genetic studies of mtDNA suggested that the emergence of the Cham was primarily mediated by cultural diffusion (Peng, et al. 2010). The other two AN groups, Ede and Giarai, have high frequencies of mtDNA haplogroups which are specific to Vietnam but absent in Taiwanese AN speakers (Duong, et al. 2018), and also have a high frequency of mtDNA but no partial MSY haplotype sharing with each other (Macholdt, et al. 2019). We find that the AN groups actually share less ancestry with Taiwan AN groups than do most other groups from Vietnam; however, Cham do share slightly more ancestry with the Taiwanese AN groups than do the Ede and Giarai, while the Ede and Giarai share slightly more ancestry with the AN-speaking Borneo and AA-speaking Htin Mal and Mlabri (figs. 4 and 6). Moreover, the admixture graph results show that the Ede and Giarai can be modeled as having exclusively AA-associated ancestry, while the Cham have ∼11% ancestry from an ancestor of the AN-speaking Atayal (fig. 8C and supplementary fig. S18). To sum up, the pattern we have observed in AN groups likely reflects the ancestors of the Cham coming from ISEA and interacting extensively with AA groups, which resulted in the Cham acquiring substantial AA-related ancestry. These interactions also led other AA groups to shift to AN languages (e.g., the Ede and Giarai). Thus, the AN-speaking groups of Vietnam do not reflect a purely cultural process for the spread of AN languages, but rather both migration and cultural diffusion. However, we should emphasize that additional sampling of Central and Southern Vietnamese ethnolinguistic groups is needed to fully document their interactions with the groups we have studied.

In the IBD results, we observe that the Vietnamese AN groups are mostly connected with neighboring AA groups and with an AN-speaking group from Borneo (fig. 6), which has been shown to have excess AA-related ancestry (Lipson, et al. 2014). We also observe strong IBD sharing between the Ede and Giarai over the entire size range of IBD blocks, which is consistent with the uniparental data for these two groups (Macholdt, et al. 2019). Additionally, the AN-speaking groups underwent population expansion around 300-450 years ago (fig. 5). A similar population expansion was inferred for the Giarai and Ede based on partial Y chromosome sequences (Macholdt, et al. 2019); the Cham were not included in this study. However, the inferred timing of population expansion based on the Y chromosome is much older (∼2,500 and ∼7,500 years ago for the Ede and Giarai, respectively), and was suggested to be possibly linked to the spread of the Dong Son culture (Macholdt, et al. 2019). Still, mtDNA genome sequences from the Giarai and Ede did not show any signal of expansion (Macholdt, et al. 2019). Given the uncertainty with dating events based on molecular genetic data, it may be that the same expansions are reflected in the autosomal and uniparental marker data. Alternatively, the uniparental markers may lack sufficient resolution to detect more recent expansions. Given that the time of expansion of AN groups based on genome-wide data is close to that of the Kinh and Muong, we suggest that these events may be linked.

### Hmong-Mien and Tai-Kadai

Both the HM and TK families are thought to have originated in what is now southern China (and possibly also northern Vietnam for the TK family), and the beginning of their separate migrations into MSEA dates to ∼2.5 kya (Edmondson and Gregerson 2007; Bellwood 2015). The TK and AN proto-languages might be related (Enfield, et al. 2011; Habu, et al. 2018), and TK groups from Thailand have been shown to be related to Austronesians based on modeling of mtDNA genome sequences (Kutanan, et al. 2018). We have also found that the AN-speaking Atayal is placed in the clade of TK groups (supplementary figs. S15, S16, and S20). The early TK, HM, ST, and AA groups are thought to have interacted in what is now southern China (Enfield, et al. 2011; Habu, et al. 2018). It has also been suggested that ancient tribes in southern China, the Baiyue, might be composed of several proto-AA, HM, and TK groups living together (Lee 2012). Compared to the AA and ST, closer interactions between the HM and TK have been shown in genetic studies using uniparental (Macholdt, et al. 2019) and insertion/deletion data (He, et al. 2019). A recent study further pointed out that Hmongic and Mienic groups from southern China demonstrate different genomic affinity to ST and TK groups, respectively (Xia, et al. 2019). We have also found that the Vietnamese HM and TK groups are closely related. Among them, the HM-speaking Dao in particular share more drift and, based on IBD sharing, have more recent interactions with nearby TK groups, especially Colao and Lachi (figs. 4 and 6). The Pathen also live close to the TK groups but share more drift and IBD blocks with the Hmong (fig. 6). This could be explained by the fact that the Hmong and Pathen speak languages that belong to the Hmongic branch of the family and thus might have a more recent common ancestor, while the Dao language belongs to the Mienic branch (Eberhard, et al. 2019). In contrast, the TK-speaking Colao share more with the HM groups, especially with the Hmong (fig. 6). The Colao and Hmong have strong interactions based on IBD sharing that cease around 650 years ago (fig. 6), which could be due to population decline in both of them after 600 years ago (fig. 5). The languages spoken by the Colao and Lachi both belong to the Kra branch of the TK family (Eberhard, et al. 2019), hence we suspect that the initial interaction was between early Kra and Mienic groups. Overall, the interactions we identify between the HM and TK groups is consistent with linguistic studies (Enfield, et al. 2011; Lee 2012; Habu, et al. 2018) and genetic studies using uniparental (Macholdt, et al. 2019) and insertion/deletions data (He, et al. 2019).

### Sino-Tibetan

The ST family originated in northern China ∼7 kya (Sagart, et al. 2019; Zhang, et al. 2019) and then started to move southward into MSEA ∼3 kya (Bellwood 2015). Compared to HM and TK groups, the ST groups form a relatively independent and isolated cluster (figs. 2, 6, and 8F). Yet, the Lolo and Phula share more drift with the HM and TK groups than do the other ST groups (fig. 4). In particular, the Lolo are modeled as sharing ancestors with the TK-speaking Dai and AN-speaking Atayal in the admixture graph analysis (fig. 8F). The Lolo and Phula live at lower elevations than the other ST groups, and the Phula live close to several HM and TK groups (fig. 1). While most of the ST groups show strong IBD sharing with each other, the Phula also share IBD blocks with the HM-speaking Hmong and the TK-speaking Boy in the recent time period (fig. 6). Although the ST-speaking Cong do not show strong shared drift with the HM and TK groups, they do share IBD blocks with the HM-speaking Hmong over the entire size range (fig. 6). This not only agrees with the genomic affinity between Hmongic and ST groups suggested recently (Xia, et al. 2019), but also indicates more recent interactions between the ST and HM groups.

## Conclusion

We have analyzed newly-generated genome-wide SNP data for the majority group Kinh plus 21 smaller ethnic groups from Vietnam. These ethnolinguistic groups speak languages that encompass the five major language families in MSEA. Our study shows extensive genetic diversity of the Vietnamese ethnolinguistic groups that is associated with heterogeneous ancestry sharing profiles in each language family. In contrast to previous studies suggesting a largely indigenous origin of the Vietnamese, we find evidence for extensive contact, over different time periods, between Vietnamese and other groups. However, the linguistic diversity is not completely in agreement with genetic diversity; in particular, the HM and TK groups in Vietnam demonstrate extensive interactions between populations speaking languages belonging to different families, while the AN groups likely reflect language shift involving AA groups. This study highlights the importance of dense sampling of ethnolinguistic groups, combined with genome-wide data from both extant and ancient sources, to gain insights into the history of an ethnolinguistically diverse region such as Vietnam.

## Materials and Methods

### Sample information

We sampled 259 male Vietnamese individuals (supplementary table S1) belonging to 22 ethnic groups that speak languages belonging to the five language families in Vietnam. Specifically, the ethnic groups consist of four Austro-Asiatic (AA) speaking groups (Khomu, Kinh, Mang, and Muong), three Austronesian (AN) speaking groups (Cham, Ede, and Giarai), three Hmong-Mien (HM) speaking groups (Dao, Hmong, and Pathen), six Sino-Tibetan (ST) speaking groups (Cong, Hanhi, Lahu, Lolo, Phula, and Sila), and six Tai-Kadai (TK) speaking groups (Boy, Colao, Lachi, Nung, Tay, and Thai). The mtDNA genome (Duong, et al. 2018; Macholdt, et al. 2019) and partial MSY sequences (Macholdt, et al. 2019) for most of these individuals, from 17 of the 22 ethnic groups, were published previously. The median of the geographic coordinates of the sampling locations per population are shown in fig. 1. The name, language affiliation, and census size of the ethnic groups included in this project were based on *the General Statistics Office of Vietnam* (www.gso.gov.vn; accessed April 2019 and the 2009 Vietnam Population and Housing census) and the Ethnologue (Eberhard, et al. 2019). All sample donors gave written informed consent, and this research received ethical clearance from the Institutional Review Board of the Institute of Genome Research, Vietnam Academy of Science and Technology (No. 4-2015/NCHG-HDDD) and from the Ethics Commission of the University of Leipzig Medical Faculty.

### Genotyping data set information

All sampled individuals were genotyped on the Affymetrix Axiom Genome-Wide Human Origins array (Patterson, et al. 2012). We kept only autosomal markers for our analyses, which contains 587,360 markers on the hg19 version of the human reference genome coordinates. In order to study ethnolinguistic history in Vietnam on a spatial-temporal scale, we merged both modern (Reich, et al. 2011; Patterson, et al. 2012; Lazaridis, et al. 2014; Qin and Stoneking 2015) and ancient (Yang, et al. 2017; Lipson, et al. 2018; McColl, et al. 2018) published data from populations within and around Mainland Southeast Asia (MSEA) (supplementary fig. S22 and supplementary table S1). Ancient samples were labeled by their excavation site and time period, with P: Paleolithic, Ho: Hoabinhian, N: Neolithic, BA: Bronze Age, IA: Iron Age, and Hi: Historical. Data merging was done by mergeit from EIGENSOFT version 7.2.1 (Patterson, et al. 2006). Positions with more than two variants or that were inconsistent between two datasets were excluded. For data genotyped on the Affymetrix 6.0 array, we first converted the genomic coordinates from hg18 to hg19 using CrossMap version 0.3.1 (Zhao, et al. 2014) and extracted the intersection of markers with our Vietnamese data set using the intersect command in bedtools version 2.25.0 (Quinlan and Hall 2010) before merging. However, incorporating data genotyped on the Affymetrix 6.0 array greatly decreased the number of informative sites due to the low number of intersecting markers (∼60,000), and we therefore only included the Affymetrix 6.0 data in population structure analyses. Similarly, incorporating ancient DNA data also greatly decreased the number of informative sites due to missing data, so we excluded the ancient samples from the phasing and identity by descent (IBD) analyses. For quality control, we first checked individual relatedness using KING version 2.1.6 (Manichaikul, et al. 2010) and removed one from each pair of individuals with 1^st^ degree of kinship. After that, we examined the global and within population missing site numbers using the missing command in PLINK version 1.90b5.2 (Purcell, et al. 2007). We removed modern individuals with more than 5% global missing data, and ancient individuals with less than 15,000 informative sites. Then, we excluded variant sites in modern samples with more than 5% global missing data, or 50% missing data within a population. We also used PLINK to perform Hardy-Weinberg equilibrium tests within populations and excluded variant sites with *p* value less than 0.00005. The number of individuals and sites for the filtered data used for different analyses is provided in supplementary table S3.

### Population structure analyses

We used principle components analysis (PCA) and ADMIXTURE version 1.3.0 (Alexander, et al. 2009) to visualize how the populations cluster. For both methods, variants were pruned beforehand for linkage disequilibrium using PLINK, excluding one variant from pairs with r^2^ > 0.4 within windows of 200 variants and a step size of 25 variants. We performed PCA by computing eigenvalues only from the less isolated modern populations and then projecting the more isolated modern populations (Mamanwa, Mlabri, Onge and Jehai) and the ancient samples, using smartpca from EIGENSOFT with “lsqproject” and “autoshrink” options. We performed heatmap visualization of downstream PCs using the pheatmap package in R version 3.6.0. For running the ADMIXTURE program, we also first estimated the allele frequency of the inferred ancestral populations (i.e. P parameter) using the less isolated modern populations and then projected the more isolated modern populations and the ancient samples with the -P option. From K = 2 to K = 15, we performed 100 replicates for each K with random seeds. Finally, we used pong version 1.4.7 (Behr, et al. 2016) to visualize the top 20 highest likelihood ADMIXTURE replicates for the major mode at each K.

### Allele sharing analyses

We used admixr version 0.7.1 (Petr, et al. 2019) to compute *f3*- and *f4*-statistics from ADMIXTOOLS version 5.1 (Patterson, et al. 2012), with significance assessed through block jackknife resampling across the genome. Outgroup *f3*-statistics of the form *f3*(X, Y; Outgroup) were used to measure the shared drift between populations X and Y since their divergence from the outgroup. We performed heatmap visualization of *f3* profiles using the pheatmap package in R. *f4*-statistics of the form f4(W, X; Y, Outgroup) were used to formally test whether W or X shares more ancestry with population Y. To avoid potential noise from ancient DNA damage patterns, we performed an additional run of *f4*-statistics using only transversions. We used Mbuti as the outgroup for all analyses; to ensure there is no excess shared ancestry between any test population and the outgroup, we also repeated the outgroup *f3*-statistics with French as the outgroup.

### Data phasing

We used SHAPEIT version 2.r904 (Delaneau, et al. 2012; Delaneau, et al. 2013; Delaneau, et al. 2014) with a reference panel and recombination map from the 1000 Genome Phase3 (Genomes Project, et al. 2015) to phase the modern samples. For the reference panel we used the East Asia and South Asia populations. To check the consistency of sites and strands between the reference panel and our data set, we ran SHAPEIT with -check option before phasing and excluded markers failing this check. For phasing, the accuracy of SHAPEIT can be increased by increasing the number of iterations and conditioning states on which haplotype estimation is based (Browning and Browning 2011). We used options --burn 10, --prune 10 and --main 30 for iteration number with 500 conditioning states, leaving other parameters as default.

### Identity by descent (IBD) analyses

We used refinedIBD (Browning and Browning 2013) to identify shared IBD blocks between each pair of individuals and homozygous-by-descent (HBD) blocks within each individual. We considered both identified IBD and HBD blocks as IBD blocks in our analyses, which have been called pairwise shared coalescence (PSC) segments in a previous study (Al-Asadi, et al. 2019). Then, we merged IBD blocks within a 0.6 cM gap and allowed only 1 inconsistent genotype between the gap and block regions using the program merge-ibd-segments from BEAGLE utilities (Browning and Browning 2007). These results were used to create four data sets based on the length of identified IBD blocks: 1 to 5 cM, 5 to 10 cM, over 10 cM, and at least 2 cM. The first three were used to compare the IBD sharing between populations in different time periods (Al-Asadi, et al. 2019), while the last one was used to investigate IBD sharing within each population (Browning and Browning 2015; Browning, et al. 2018). To summarize the IBD sharing, we summed up the total number and length of IBD blocks for each individual pair and calculated the population median and mean for each data set. We used the network approach in Cytoscape version 3.7.1 (Shannon, et al. 2003) to visualize the results, and kept the pairs with at least 2 shared blocks (4 for the range of 1 to 5 cM) to reduce noise and false positives. To estimate effective population size, we ran IBDNe (Browning and Browning 2015; Browning, et al. 2018) using shared blocks of at least 2cM within each population, and only extracted the estimated population size numbers within 50 generations ago, as previously suggested for SNP array data (Browning and Browning 2015). A generation time of 30 years (Fenner 2005) was used to convert generations to years.

### Admixture graph analyses

We used admixture graphs to model population histories that fit the genomic data. We separated our Vietnamese data by language family and modeled the admixture graph, together with related source populations, for each family. We first modeled a global admixture graph with the related present-day source populations, ancient samples, and all the Vietnamese groups. These present-day source populations were chosen based on excess ancestry sharing in the *f4* analyses. Only the ancient samples with less than 65% missing data were used here in order to have at least 20,000 SNPs for the model estimation. As the ancient samples are not closely related to the Vietnamese groups in the global admixture analysis, and their inclusion decreases the number of SNPs while increasing the complexity of the modeling, we decided to use only the present-day source populations for dissecting the Vietnamese admixture graph. We first modeled an admixture graph with only the related modern source populations, which we call the backbone populations. For each language family and the backbone populations, we pruned the SNPs as we did in the population structure analyses and calculated allele frequencies with PLINK. Using the covariance of the allele frequency profiles as input, we first ran TreeMix version 1.12 (Pickrell and Pritchard 2012) with 0 to 3 migration events and 10 independent runs, and selected the topology with the highest likelihood for further investigation. We also checked and confirmed that the likelihood and topologies of these 10 runs are mostly similar, which indicates that the model estimation has reached convergence. Next, we used AdmixtureBayes (Nielsen 2018) to estimate the top 10 posterior admixture graphs, based on the covariance of the allele frequency profiles. When more populations are added to the model, more steps will be needed for the MCMC to converge. We hence kept the maximum number of population to 11, for which the model can converge and finish in a reasonable time. To do so, we selected suitable combinations of source populations for each language family, based on the topology showing the lowest standard error in the TreeMix residual plots with 3 migration events. We used 300,000 MCMC steps for each AdmixtureBayes run with stop criteria stopping the run if the summaries of effective sample size are all above 200. We then used the estimated graphs as input for qpGraph from ADMIXTOOLS to test the goodness of fit of the graphs. We accepted the graph as a good fit when the absolute value of the Z-score of the worst *f4* statistic output by qpGraph was less than 3. For the cases where we failed to find a fit, we adjusted the source populations and excluded some highly admixed populations such as Kinh, Cong, and Colao, based on the *f4* outliers output by qpGraph. Then, we used the --subnodes option in AdmixtureBayes to calculate the posterior of the adjusted subsets and tested the results again in qpGraph. We iterated these procedures until we were able to fit graphs for all of the five language families as well as only the source populations. We ran qpGraph with parameters outpop: NULL, useallsnps: YES, blgsize: 0.05, forcezmode: YES, lspmode: YES, diag: .0001, bigiter: 6, hires: YES, and lambdascale: 1.

## Data Availability

To comply with the informed consent under which the samples were obtained, we make the data available upon request by asking the person requesting the data to agree in writing to the following restrictions: 1) The data will only be used for studies of population history; 2) the data will not be used for medical or disease-related studies, or for studies of natural selection; 3) the data will not be distributed to anyone else; 4) the data will not be used for any commercial purposes; and 5) no attempt will be made to identify any of the sample donors.

## Acknowledgments

This research was funded by the Ministry of Science and Technology, Vietnam (DTDL.CN-05/15) and by the Max Planck Society. We thank the sample donors for contributing to this research. We thank David Reich for assistance with genotyping. We thank Enrico Macholdt, Irina Pugach, Wibhu Kutanan, and Ben Peter for helpful discussion and for assistance with analyses. We thank the Multimedia Department of the Max Planck Institute for Evolutionary Anthropology for assistance with figures. B.P. acknowledges the LABEX ASLAN (ANR-10-LABX-0081) of Université de Lyon for its financial support within the program “Investissements d’Avenir” (ANR-11-IDEX-0007) of the French government operated by the National Research Agency (ANR).

## Author Contributions

M.S., B.P., and N.V.H. conceived the study. M.S. and N.V.H. funded the study. N.T.D., N.D.T., N.V.P., and N.V.H. collected the samples. N.T.D., N.D.T., and N.V.P. performed the laboratory work. D.L. analyzed the data with input from M.S., B.P., and N.V.H. D.L., M.S., and N.V.H. wrote the manuscript with input from all co-authors.

