## Supplemental captions for "Extensive ethnolinguistic diversity in Vietnam reflects multiple sources of genetic diversity"

### Supplementary figures (see 03.FigS1-22.pdf)

**Fig. S1. PCA colored by countries.** Black dots represent the ancient samples; populations are numbered according to supplementary table S1.

**Fig. S2. Heatmap visualization of PC1 to PC10.** Eigenvectors were normalized to range from 0 to 1. Each column stands for a single individual, with individuals grouped by ethnic group (labels at the bottom). The color bar denotes the language families.

**Fig. S3. Cross-validation error of ADMIXTURE runs for K= 2 to K = 15, based on 100 runs for each K value.**

**Fig. S4. ADMIXTURE results for K = 2 to K = 15.** The ancient samples are also shown separately in supplementary fig. S5.

**Fig. S5. ADMIXTURE results for the ancient samples for K = 2 to K = 15.**

**Fig. S6. Map visualization of outgroup *f3* profiles of Vietnamese ethnolinguistic groups, compared with other Vietnamese groups.** Outgroup *f3* values were normalized to range from 0 to 1. Each panel depicts an ethnolinguistic group, with the cross denoting the sample location. Labels indicate groups that are among the top 1% *f3* values with the panel group. Groups with same sample locations were jittered for visibility. The panels are colored according to language family. The Mbuti were used as the outgroup.

**Fig. S7. Map visualization of outgroup *f3* profiles of Vietnamese ethnolinguistic groups compared with nearby modern populations (Mbuti as outgroup).** Outgroup *f3* values were normalized to range from 0 to 1. Each panel depicts an ethnolinguistic group, with the cross denoting the sample location. The panels are colored according to language family. Population labels indicate groups that are among the top 1% *f3* values with the panel group.

**Fig. S8. Map visualization of outgroup *f3* profile of Vietnamese ethnolinguistic groups compared with nearby modern populations (French as outgroup).** Outgroup *f3* values were normalized to range from 0 to 1. Each panel depicts an ethnolinguistic group, with the cross denoting the sample location. The panels are colored according to language family. Population labels indicate groups that are among the top 1% *f3* values with the panel group.

**Fig. S9. Map visualization of outgroup *f3* profile of Vietnamese ethnolinguistic groups compared with the ancient samples.** Outgroup *f3* values were normalized to range from 0 to 1. Each panel depicts an ethnolinguistic group, with the cross denoting the sample location. Population labels indicate groups that are among the top 1% *f3* values with the panel group. Groups with same sample locations were jittered for visibility. The panels are colored according to language family. The Mbuti were used as the outgroup.

**Fig. S10. IBD sharing within each Vietnamese ethnolinguistic group.** The Y axis and X axis indicate the mean number and summed length of IBD blocks within each group, respectively. Ethnic groups are colored according to language family.

**Fig. S11. *f4* statistics comparing Vietnamese groups to the ancient samples.** Z-scores are for *f4*(W, southern Han Chinese; Y, Mbuti), where W is the ancient sample (panel labels) and Y is the Vietnamese group (Y axis label). Ancient samples are ordered according to periods. The vertical grey lines denote 0.

**Fig. S12. *f4* statistics comparing the ancient samples to the TK, HM, and ST groups and AA and AN groups.** Z-scores are for *f4*(TK, HM, and ST groups, AA and AN groups; Ancient samples, Mbuti), where a significant (|Z| > 3) positive value indicates excess ancestry sharing with the TK, HM, and ST groups (Dai, Miao, Chinese Lahu, southern Han Chinese, and northern Han Chinese) while a significant negative value indicates excess ancestry sharing with the AA and AN groups (Mlabri, Htin Mal, Atayal, Borneo, Ami, and Mamanwa). Ancient samples are ordered according to periods. The vertical grey lines denote 0.

**Fig. S13. *f4* statistics comparing Vietnamese groups to the ancient samples, using only transversions.** Z-scores are for *f4*(W, southern Han Chinese; Y, Mbuti), where W is the ancient sample (panel labels) and Y is the Vietnamese group (Y axis label). Ancient samples are ordered according to periods. The vertical grey lines denote 0.

**Fig. S14. *f4* statistics comparing the ancient samples to the TK, HM, and ST groups and AA and AN groups, using only transversions.** Z-scores are for *f4*(TK, HM, and ST groups, AA and AN groups; Ancient samples, Mbuti), where significant (|Z| > 3) positive values indicate excess ancestry sharing with the TK, HM, and ST groups (Dai, Miao, Chinese Lahu, southern Han Chinese, and northern Han Chinese) while significant negative values indicate excess ancestry sharing with the AA and AN groups (Mlabri, Htin Mal, Atayal, Borneo, Ami, and Mamanwa). Ancient samples are ordered according to periods. The vertical grey lines denote 0.

**Fig. S15. Global TreeMix results with 0 to 3 migrations.** The phylogenetic tree is on the left and the corresponding residual plot on the right. The Vietnamese AA, AN, HM, TK, ST groups are colored in purple, brown, yellow, red, and green, respectively.

**Fig. S16. TreeMix results for the backbone populations with 0 to 3 migrations.** The phylogenetic tree is on the left and the corresponding residual plot on the right.

**Fig. S17. TreeMix results for the Vietnamese AA groups with 0 to 3 migrations.** The phylogenetic tree is on the left and the corresponding residual plot on the right. The Vietnamese AA groups are colored in purple.

**Fig. S18. TreeMix results for the Vietnamese AN groups with 0 to 3 migrations.** The phylogenetic tree is on the left and the corresponding residual plot on the right. The Vietnamese AN groups are colored in brown.

**Fig. S19. TreeMix results for the Vietnamese HM groups with 0 to 3 migrations.** The phylogenetic tree is on the left and the corresponding residual plot on the right. The Vietnamese HM groups are colored in yellow.

**Fig. S20. TreeMix results for the Vietnamese TK groups with 0 to 3 migrations.** The phylogenetic tree is on the left and the corresponding residual plot on the right. The Vietnamese TK groups are colored in red.

**Fig. S21. TreeMix results for the Vietnamese ST groups with 0 to 3 migrations.** The phylogenetic tree is on the left and the corresponding residual plot on the right. The Vietnamese ST groups are colored in green.

**Fig. S22. Sample information map.** Map with comparative data used in this study. The crosses denote samples genotyped on the Affymetrix 6.0 platform. (A) Samples colored according to language family. (B) Samples colored according to time period.

### Supplementary tables (see 04.TableS1-3.xlsx)

**Table S1. Meta information for each sample.** Ancient samples were labeled by their time period and excavation site. Column “Population#” refers to the numbers in the PC plots. “Ancient” in Platform column means that samples were generated through ancient DNA sequencing procedures instead of SNP array (see the references in Reference column for details). Han_SC: southern Han Chinese, Han_NC: northern Han Chinese, Lahu_C: Chinese Lahu, and Thai_T: Thai from Thailand.

**Table S2. Meta information for each analysis.**

**Table S3. Source proportions in the Vietnamese groups and ancient samples for K = 6.** P: Paleolithic, Ho: Hoabinhian, N: Neolithic, BA: Bronze Age, IA: Iron Age, and Hi: Historical.
