## Supplemental Fig. S1-22 for "Extensive ethnolinguistic diversity in Vietnam reflects multiple sources of genetic diversity"

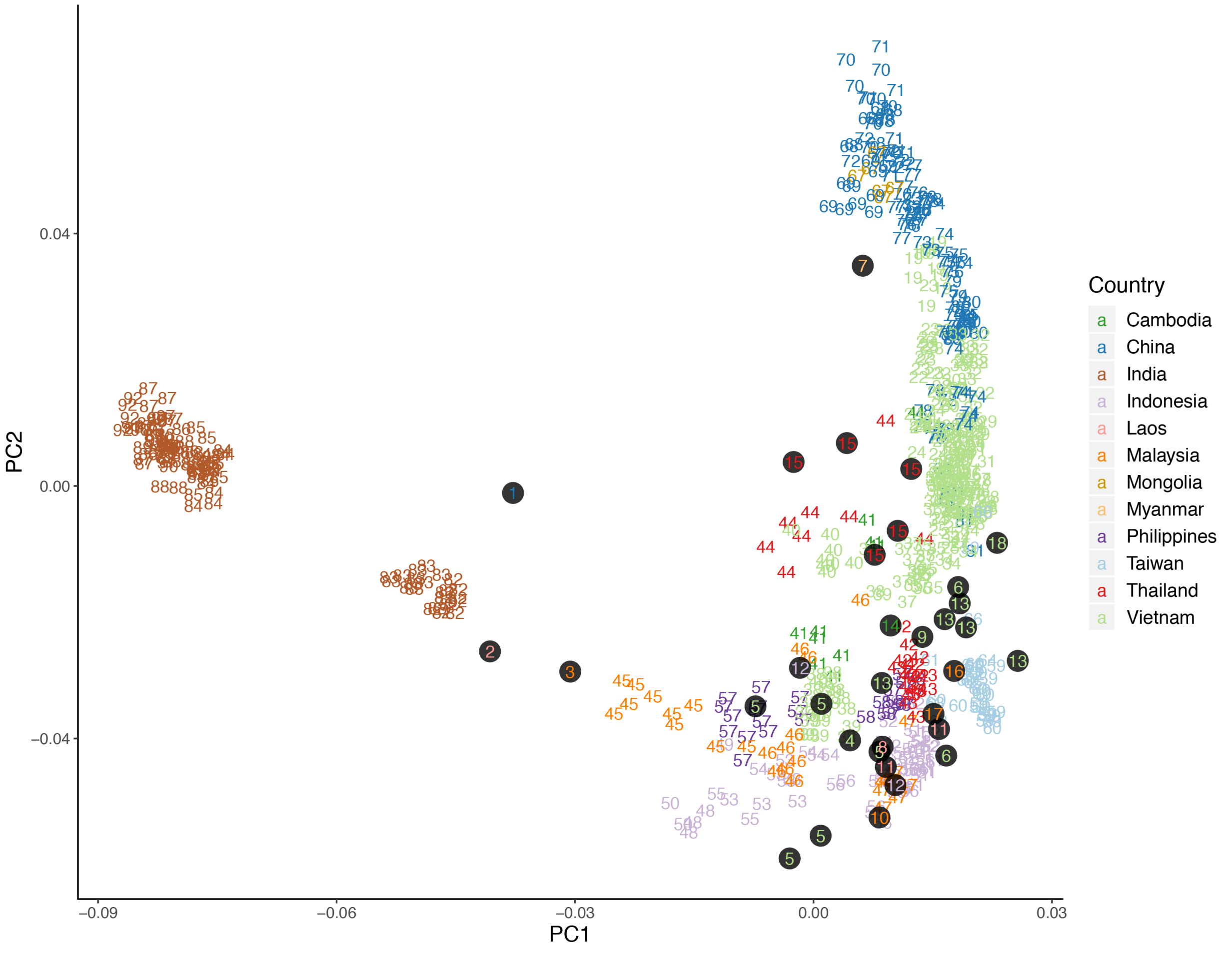


**Fig. S1. PCA colored by countries.**

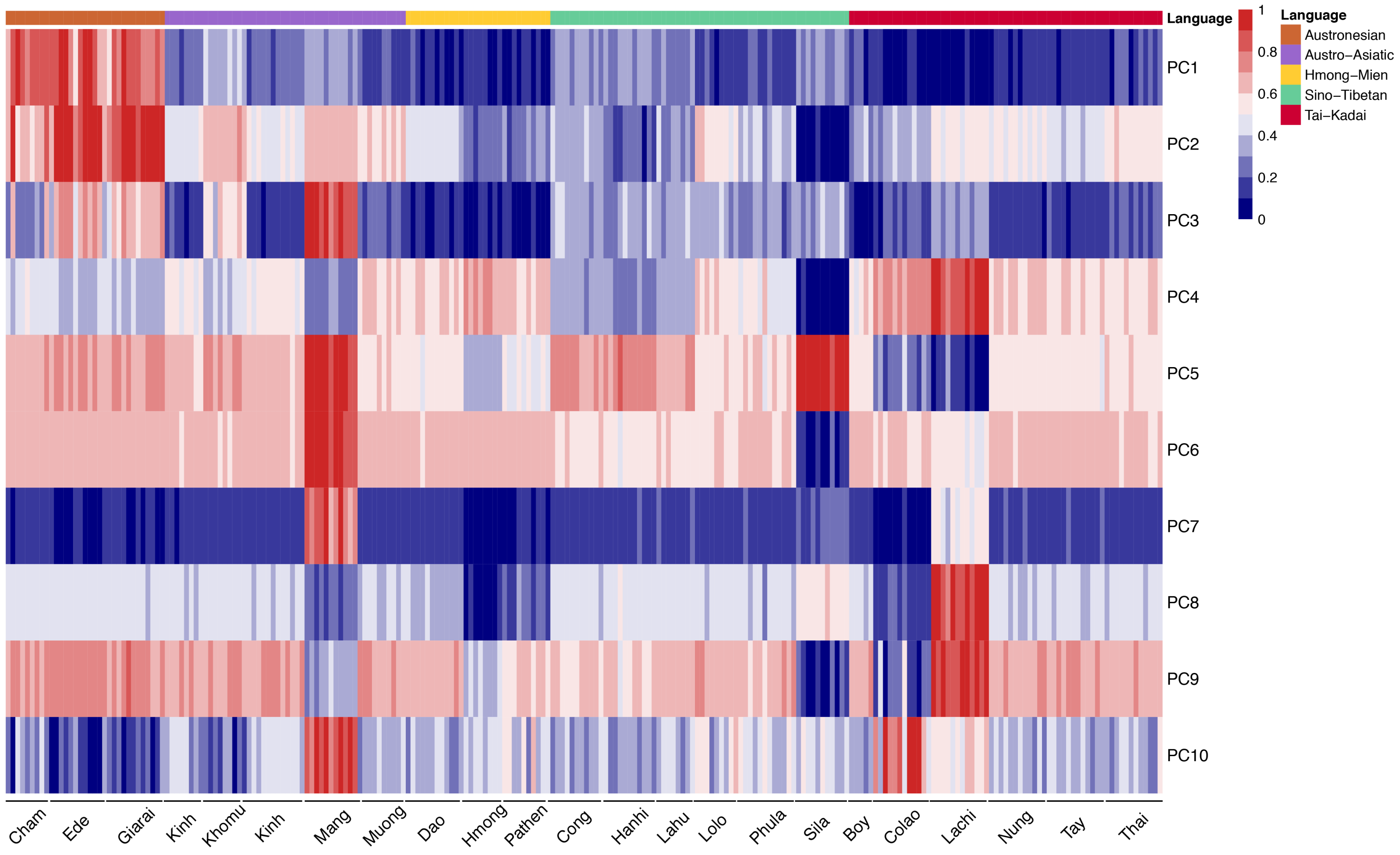


**Fig. S2. Heatmap visualization of PC1 to PC10.**

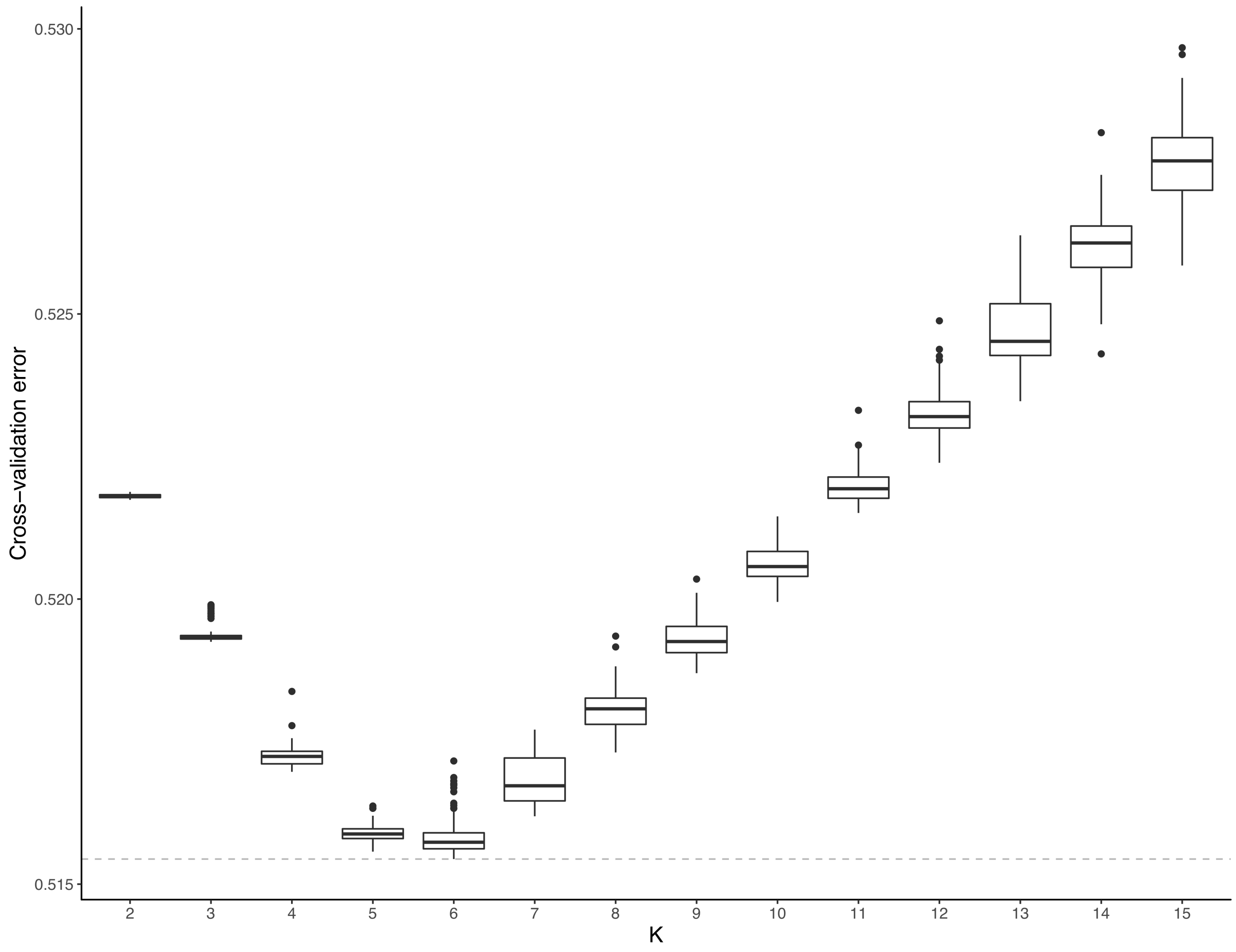


**Fig. S3. Cross-validation error of ADMIXTURE runs for K= 2 to K = 15, based on 100 runs for each K value.**

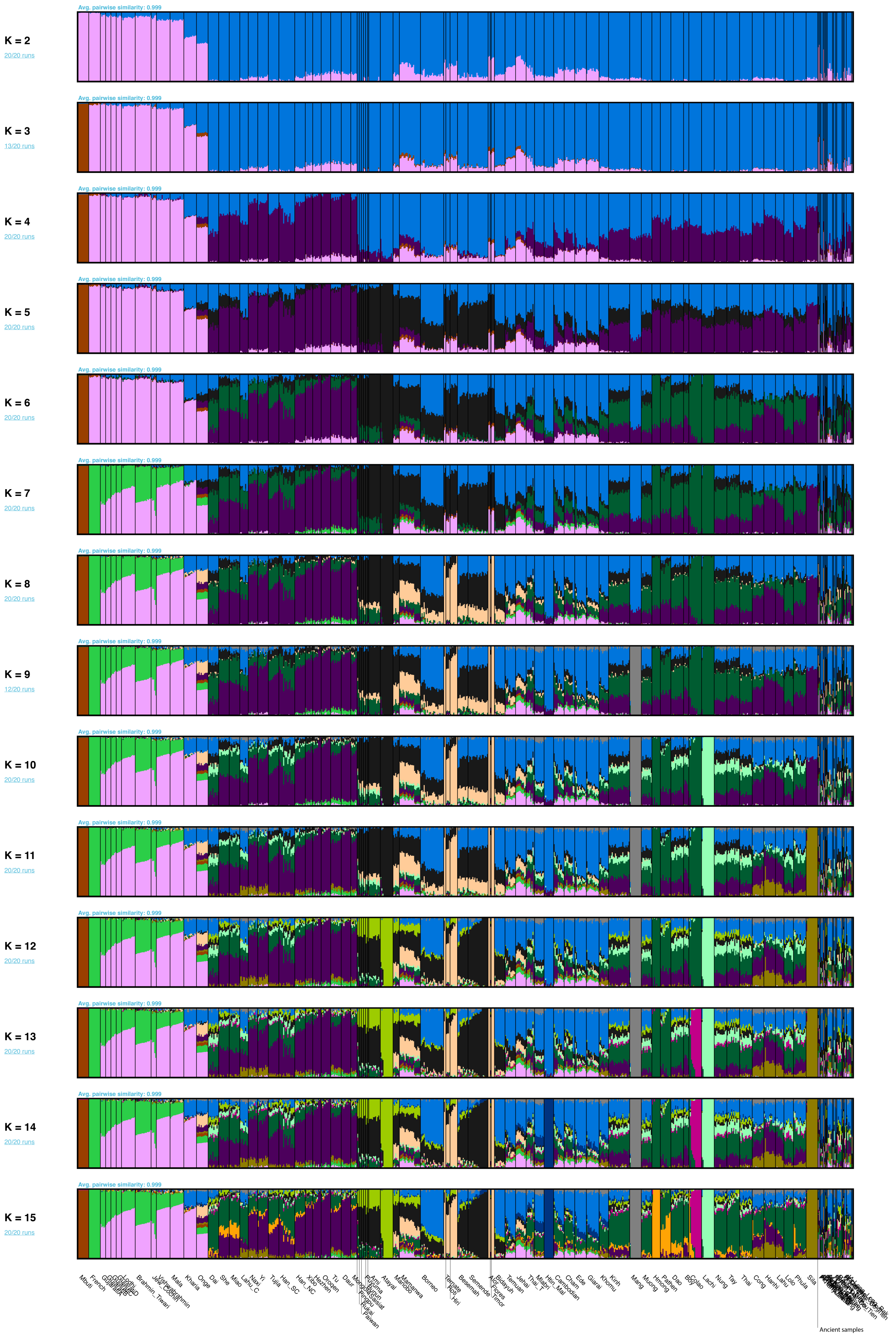

**Fig. S4. ADMIXTURE results for K = 2 to K = 15.**

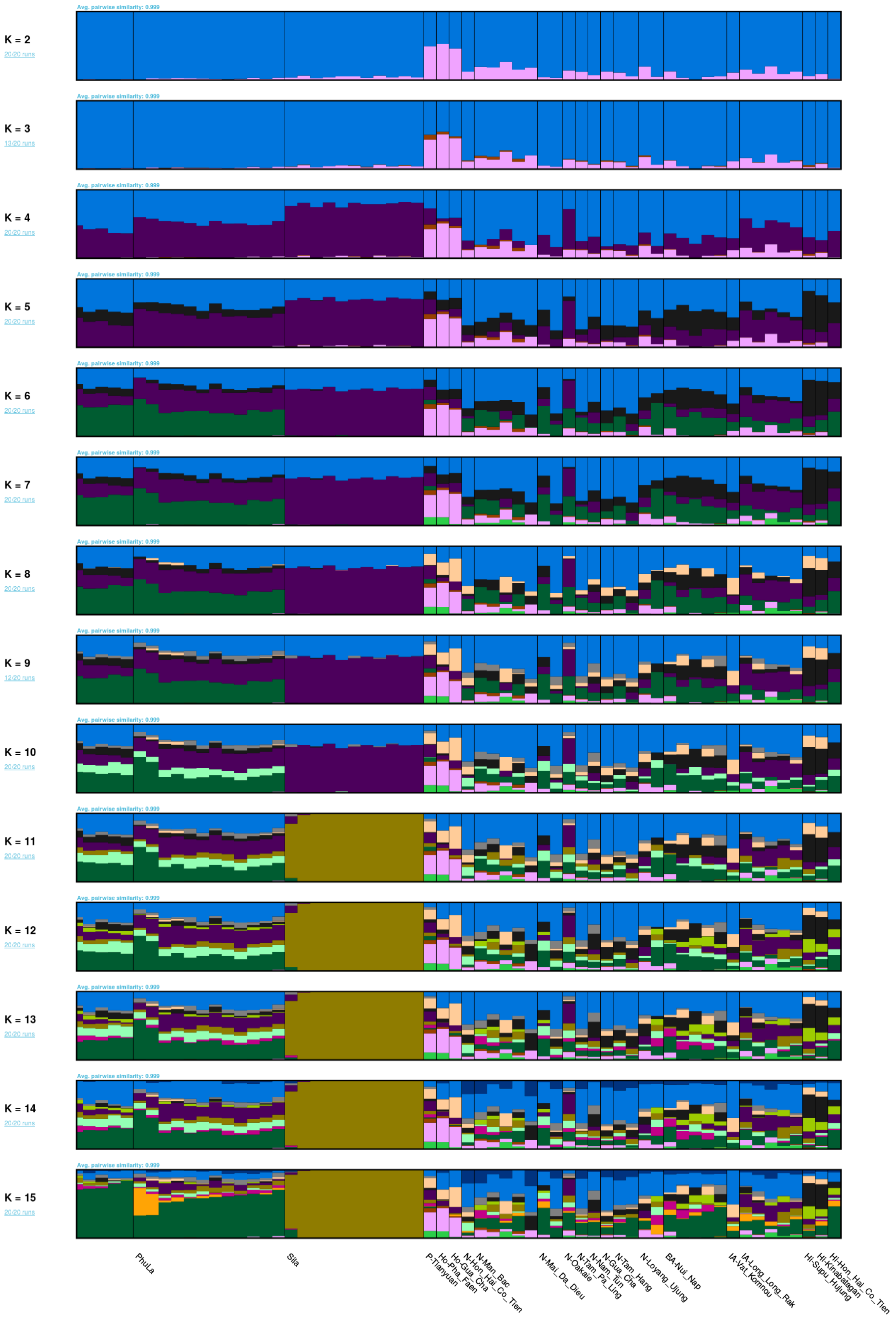


**Fig. S5. ADMIXTURE results for the ancient samples for K = 2 to K = 15.**

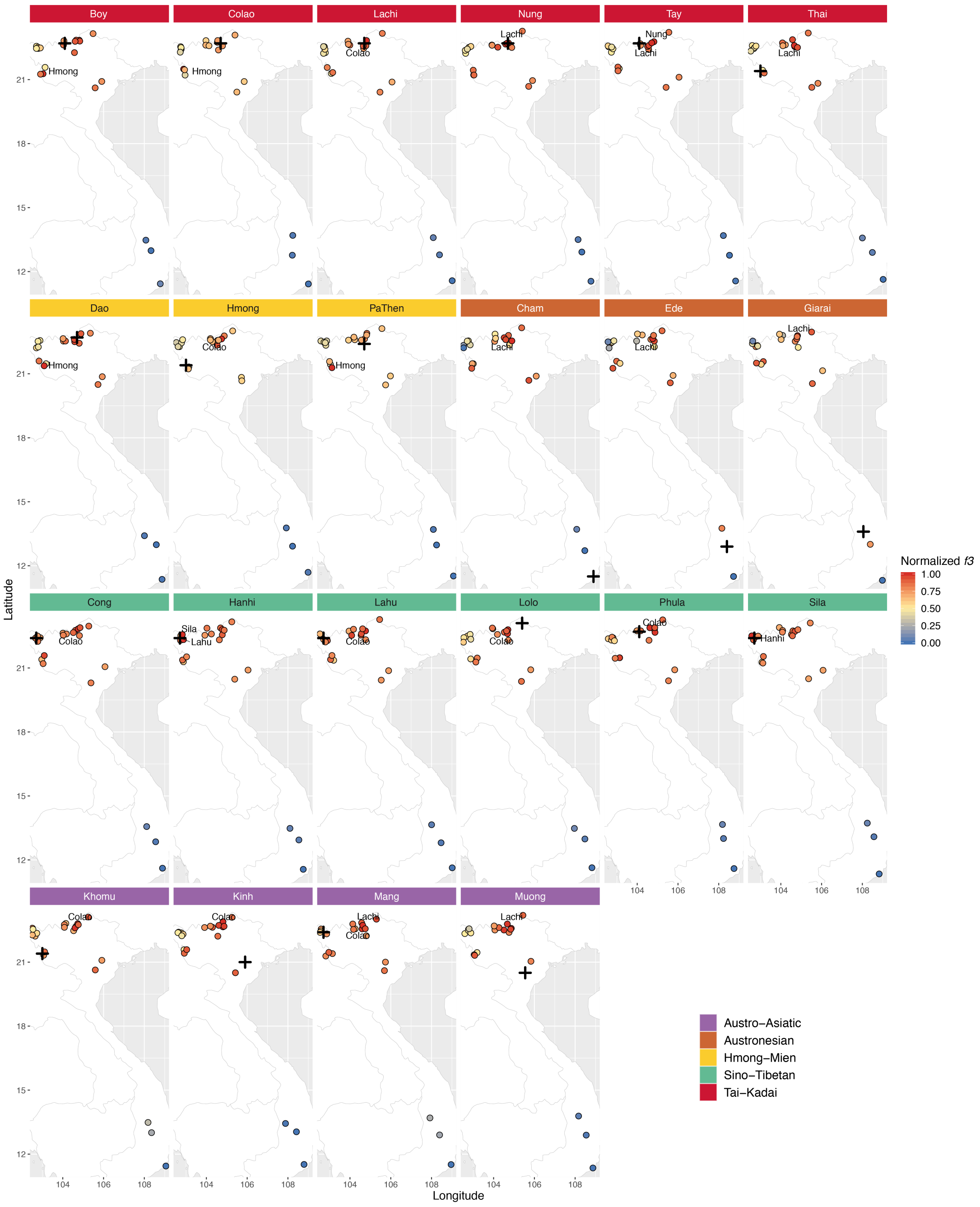


**Fig. S6. Map visualization of outgroup *f3* profiles of Vietnamese ethnolinguistic groups, compared with other Vietnamese groups.**

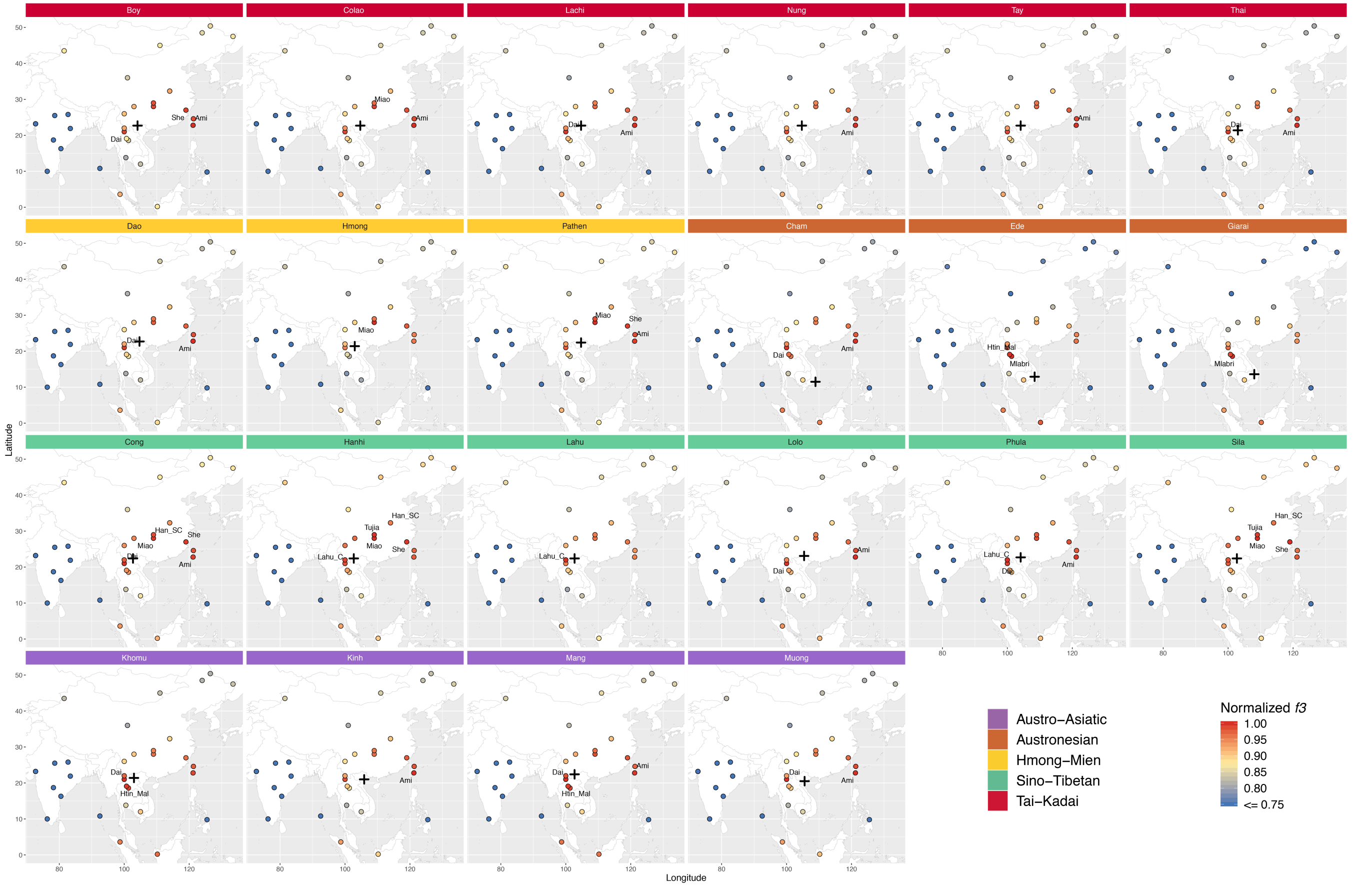


**Fig. S7. Map visualization of outgroup *f3* profiles of Vietnamese ethnolinguistic groups compared with nearby modern populations (Mbuti as outgroup).**

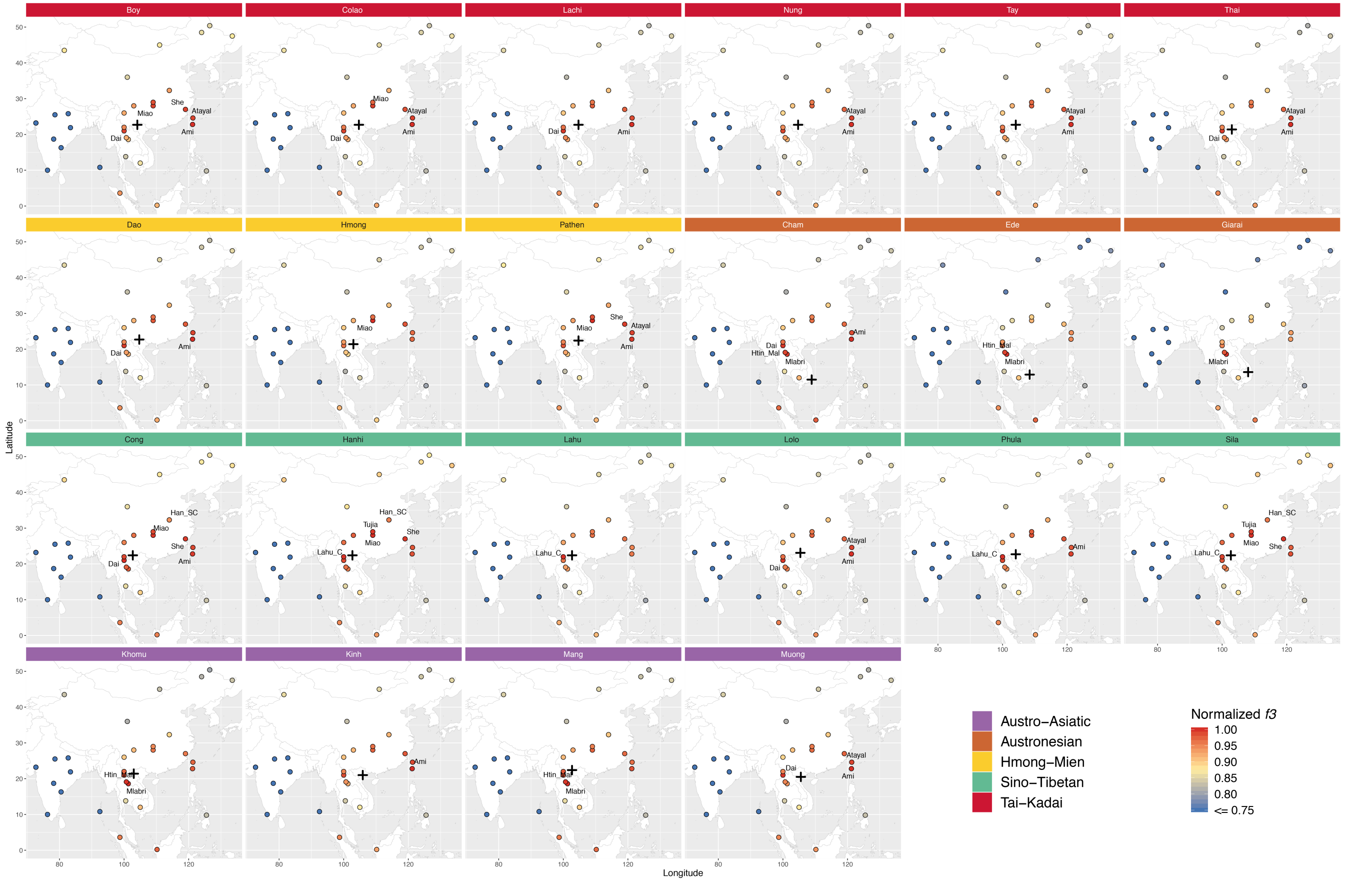


**Fig. S8. Map visualization of outgroup *f3* profile of Vietnamese ethnolinguistic groups compared with nearby modern populations (French as outgroup).**

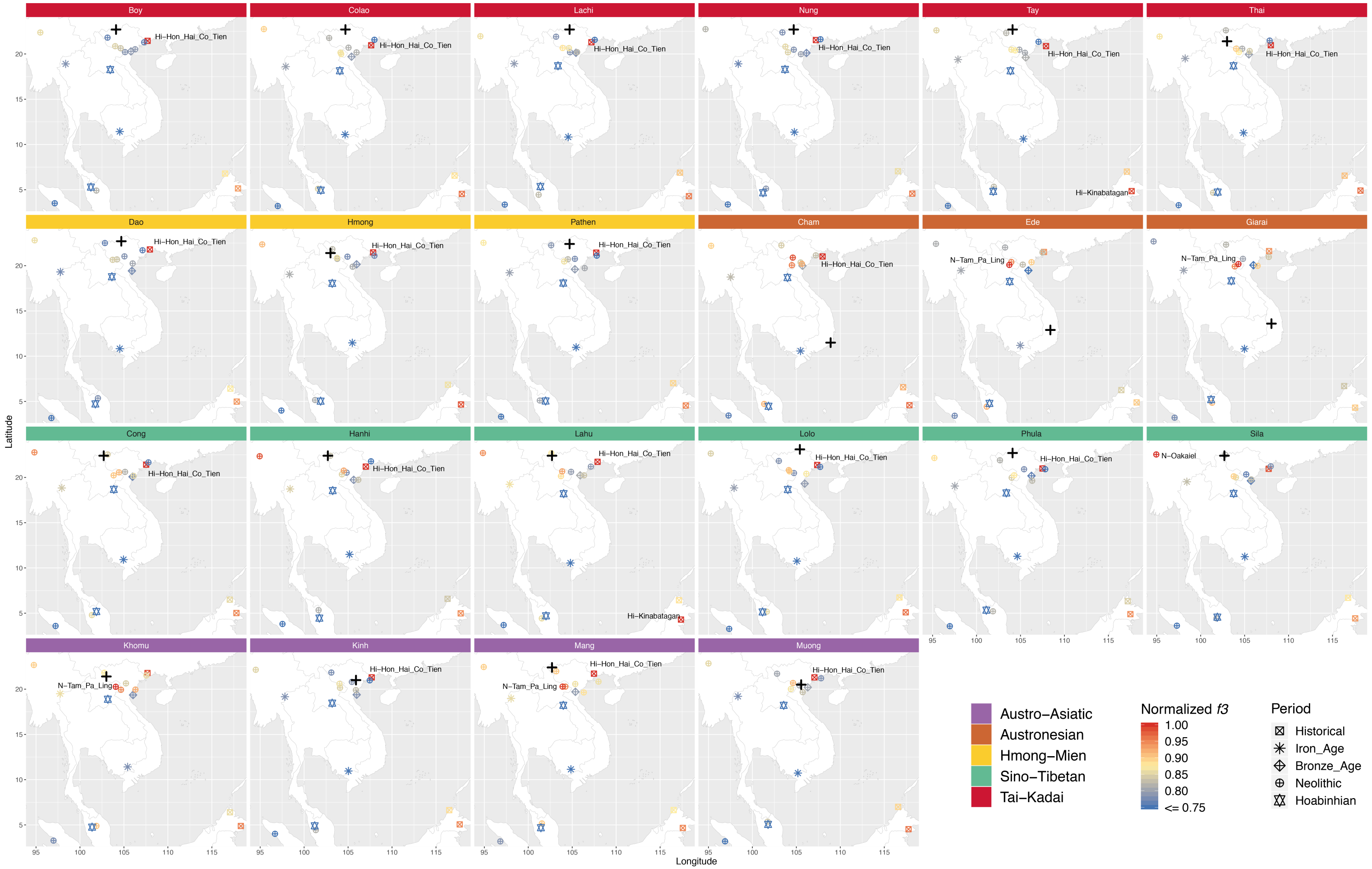


**Fig. S9. Map visualization of outgroup *f3* profile of Vietnamese ethnolinguistic groups compared with the ancient samples.**

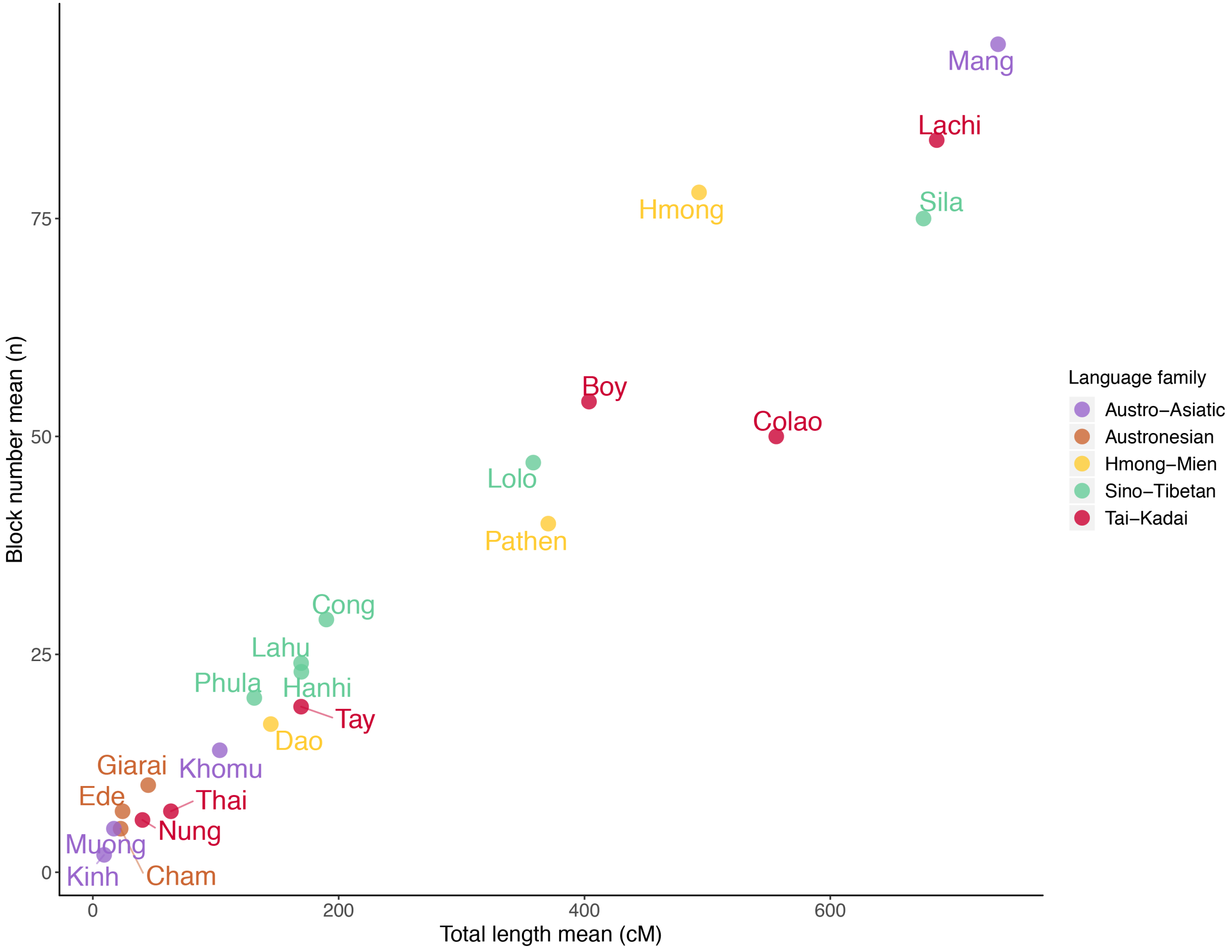


**Fig. S10. IBD sharing within each Vietnamese ethnolinguistic group.**

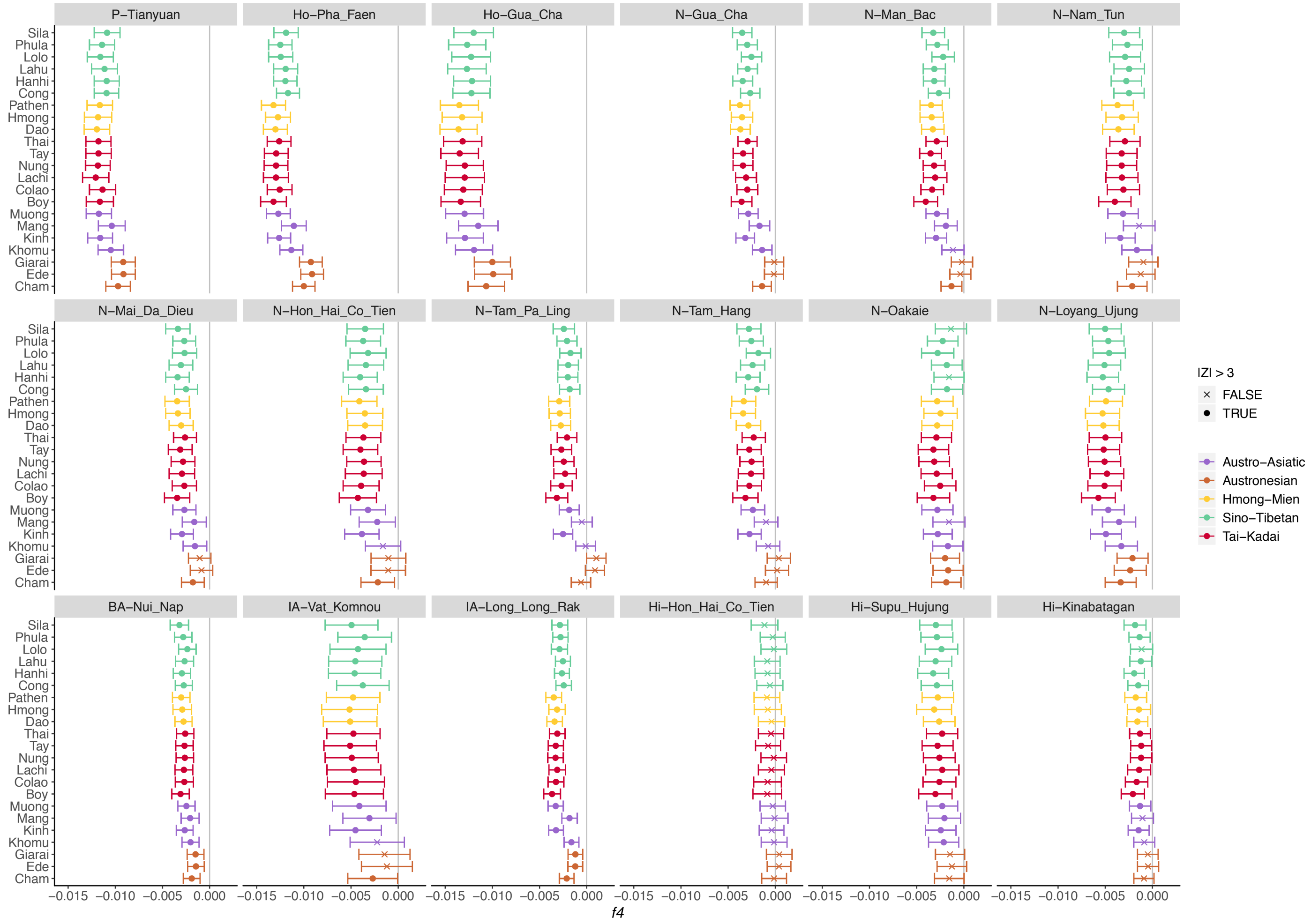


**Fig. S11.** ***f4*** **statistics comparing Vietnamese groups to the ancient samples.**

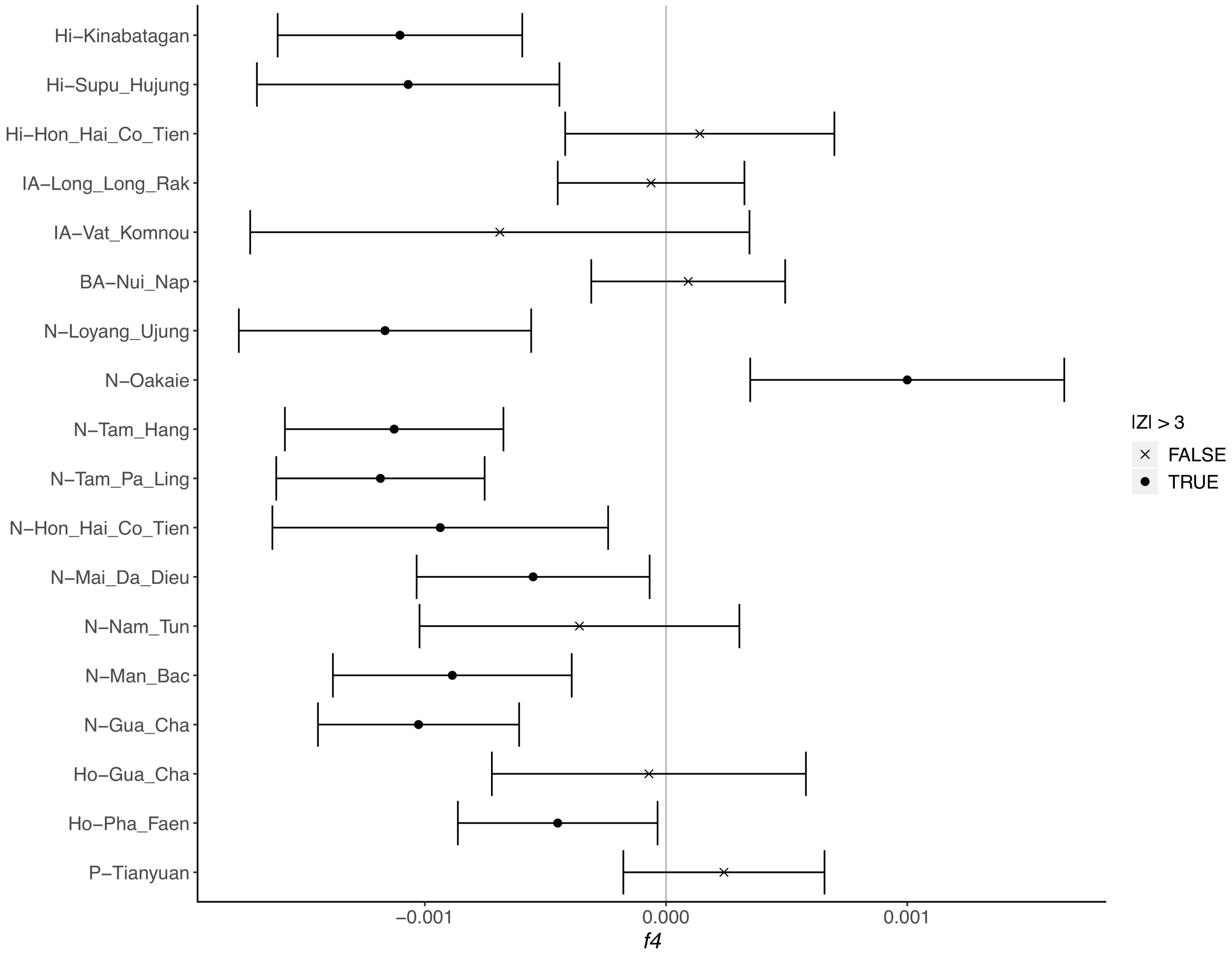


**Fig. S12.** ***f4*** **statistics comparing the ancient samples to the TK, HM, and ST groups and AA and AN groups.**

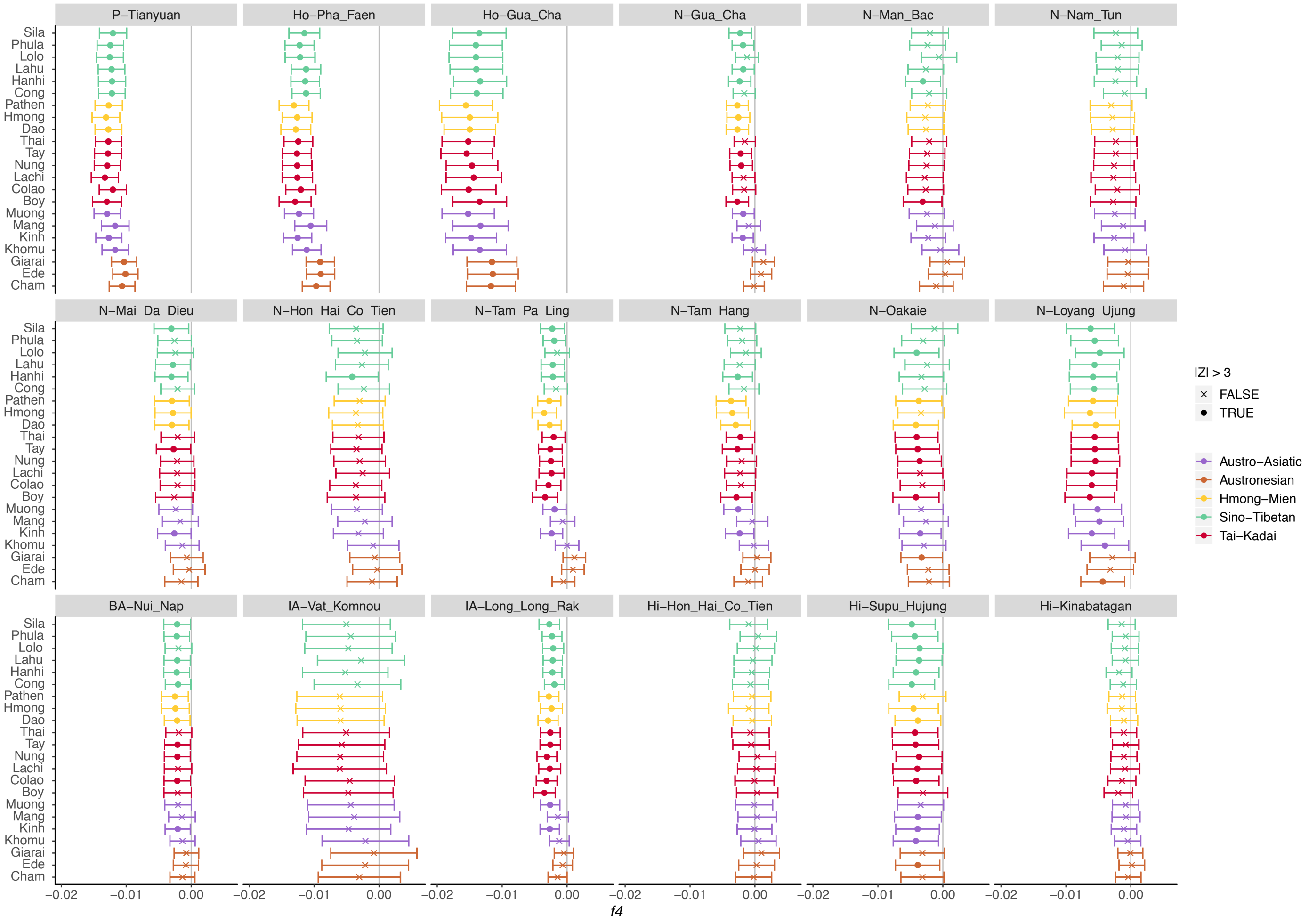


**Fig. S13.** ***f4*** **statistics comparing Vietnamese groups to the ancient samples, using only transversions.**

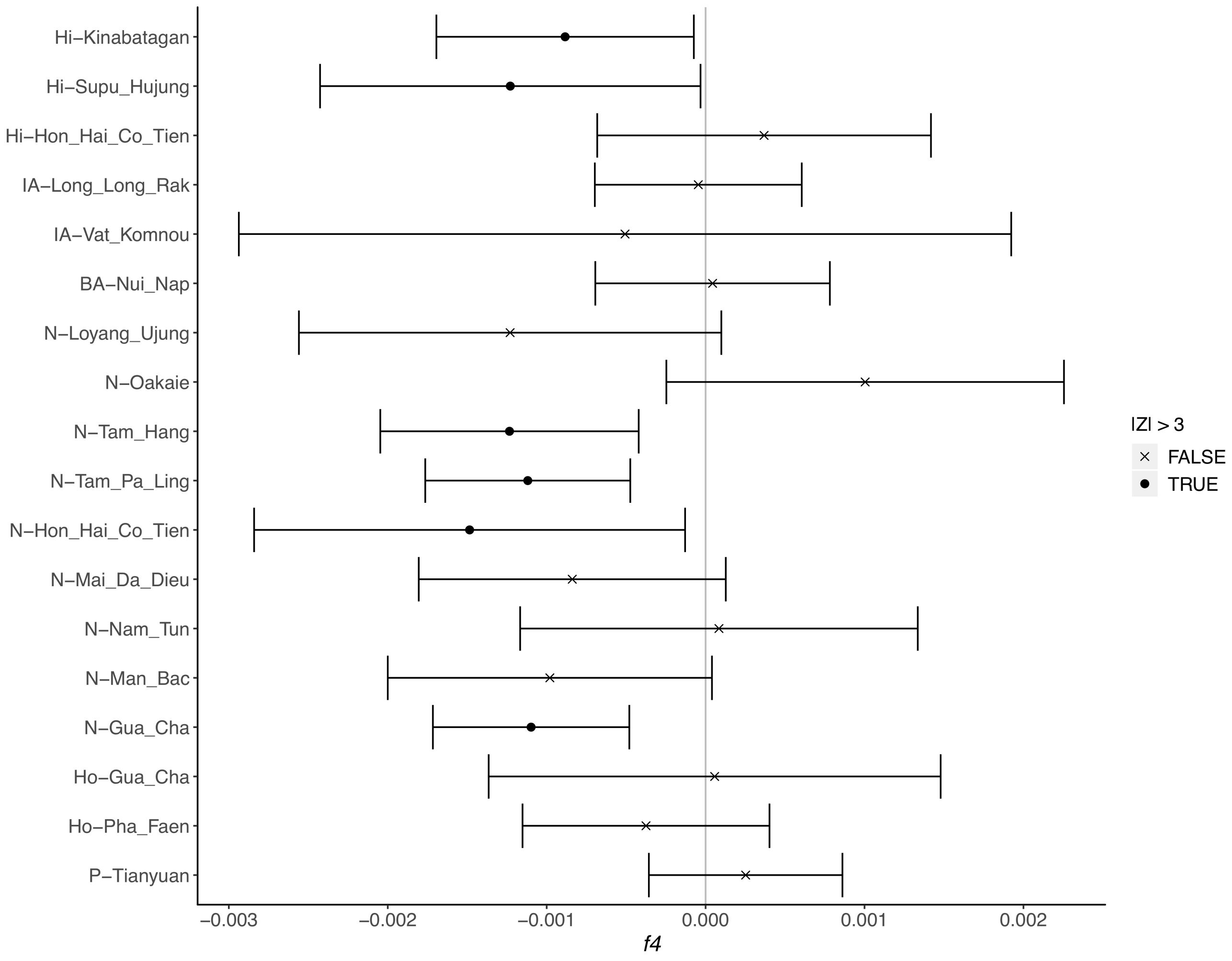


**Fig. S14.** ***f4*** **statistics comparing the ancient samples to the TK, HM, and ST groups and AA and AN groups, using only transversions.**

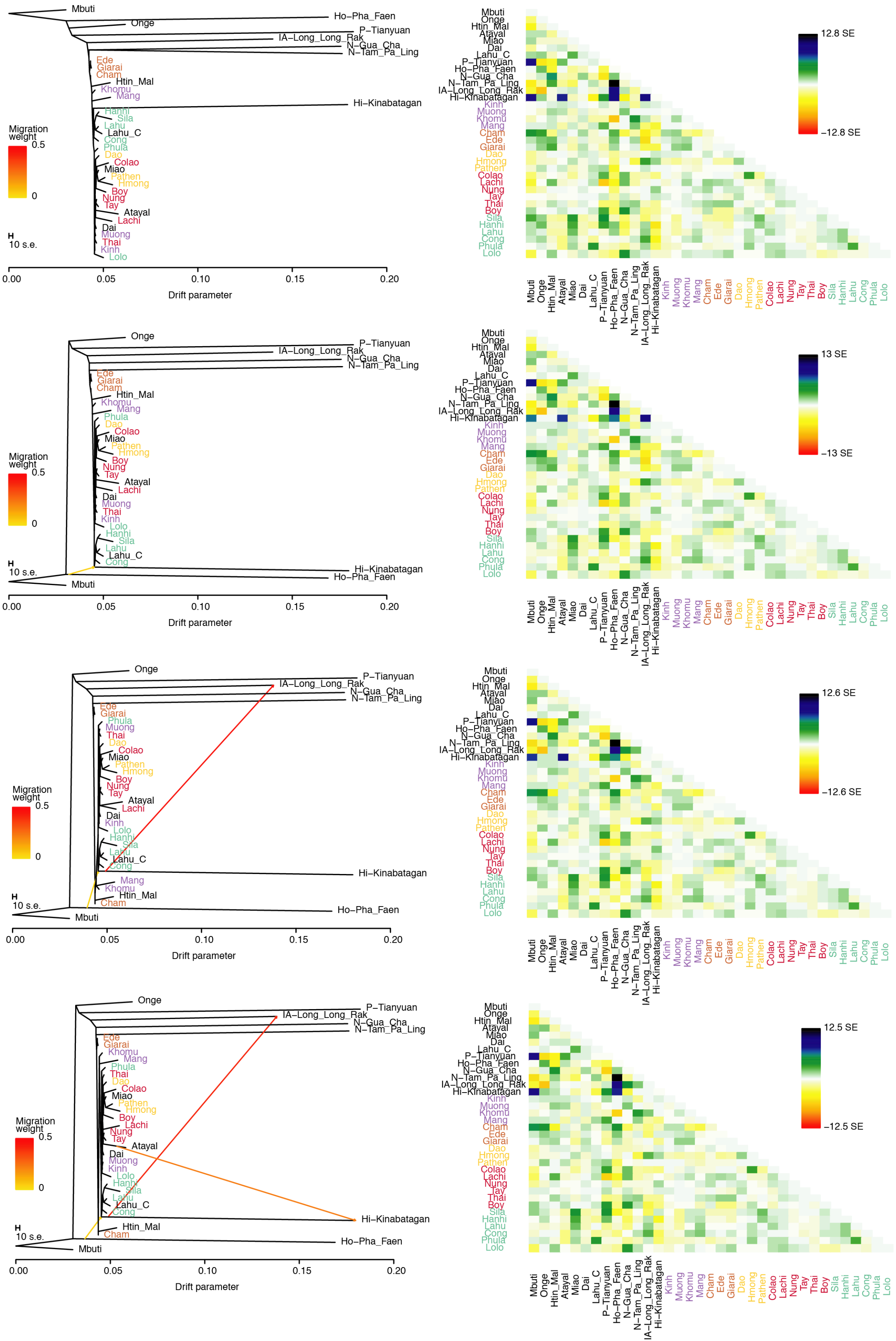


**Fig. S15. Global TreeMix results with 0 to 3 migrations.**

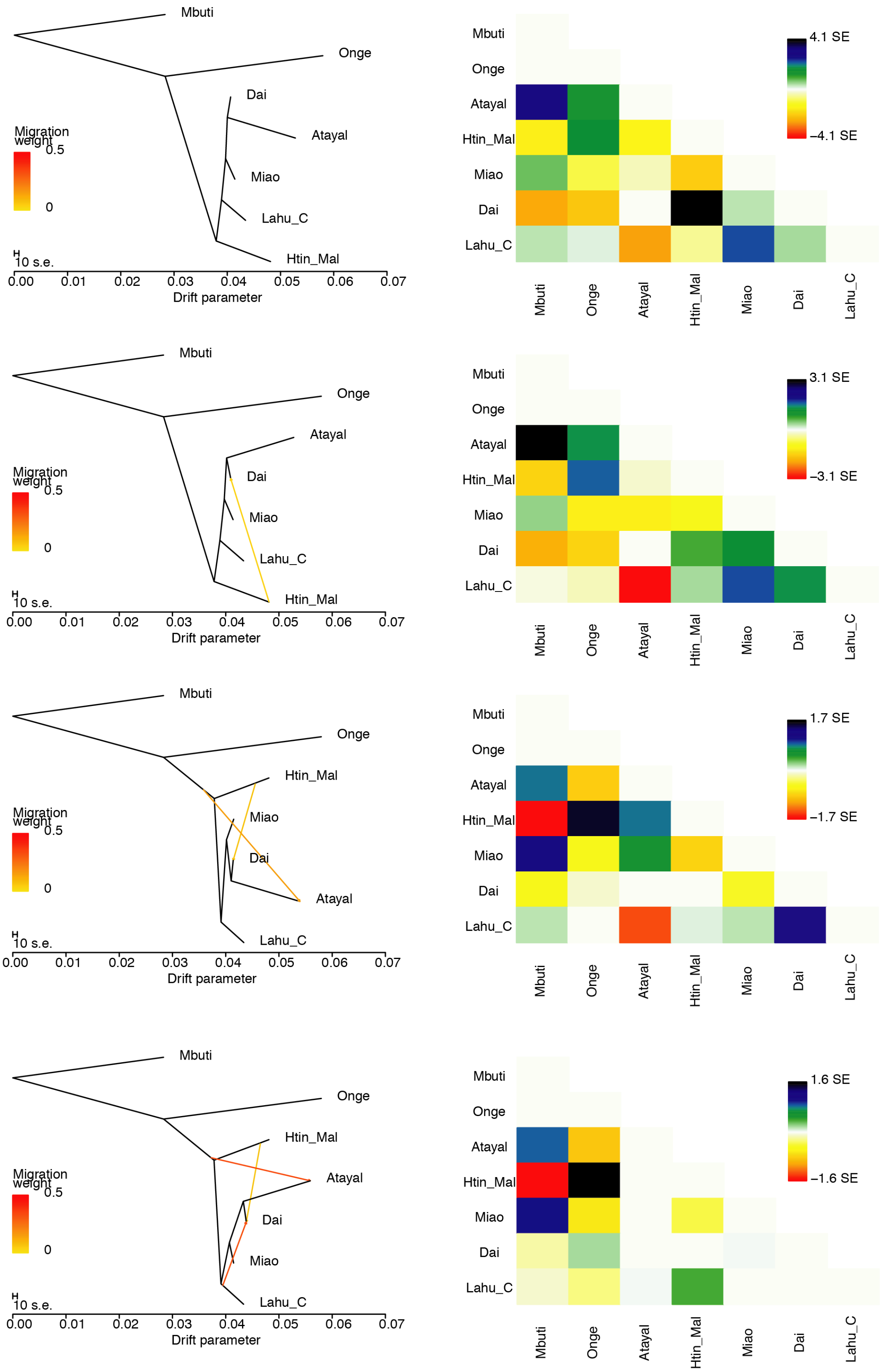


**Fig. S16. TreeMix results for the backbone populations with 0 to 3 migrations.**

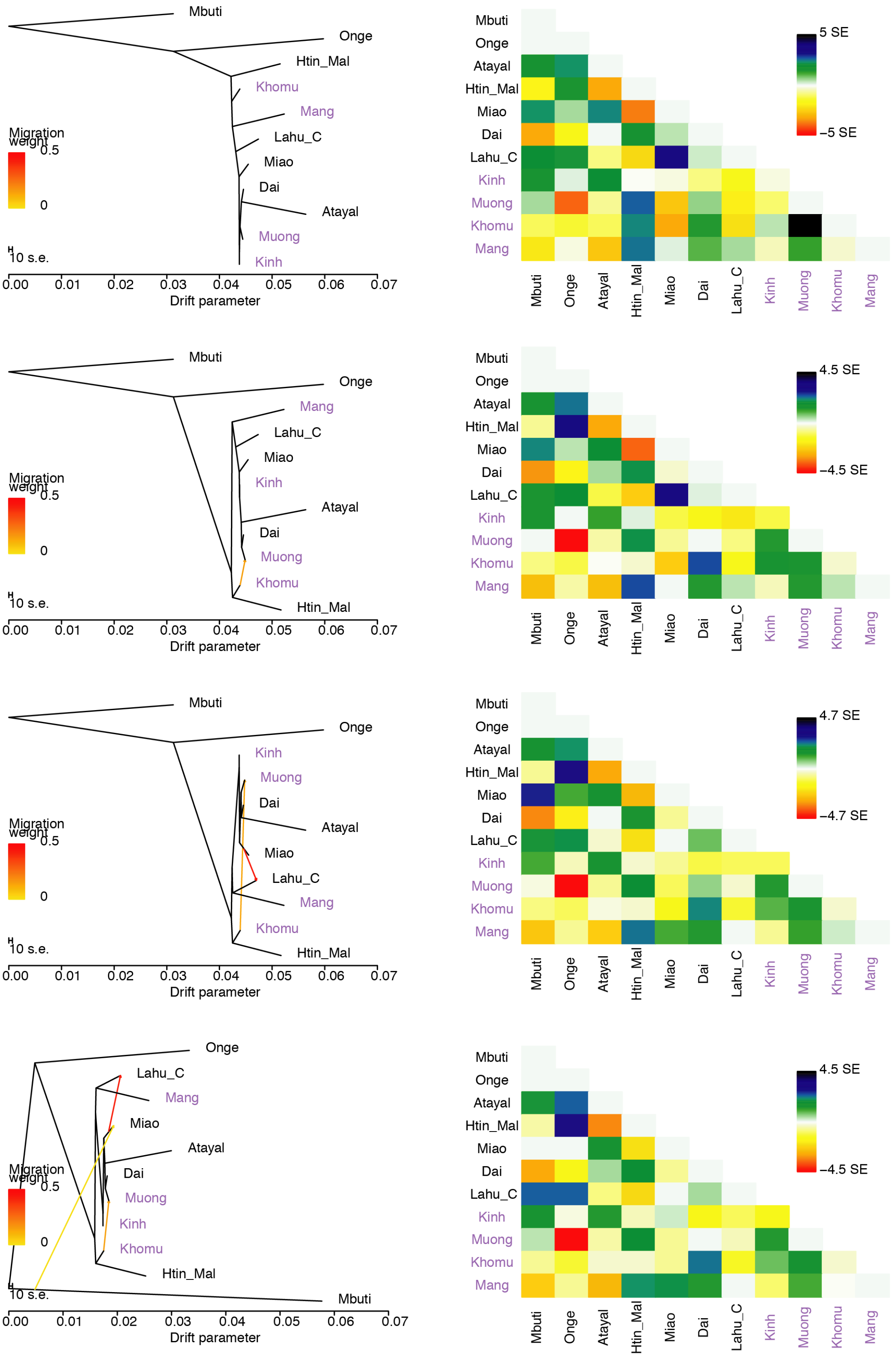


**Fig. S17. TreeMix results for the Vietnamese AA groups with 0 to 3 migrations.**

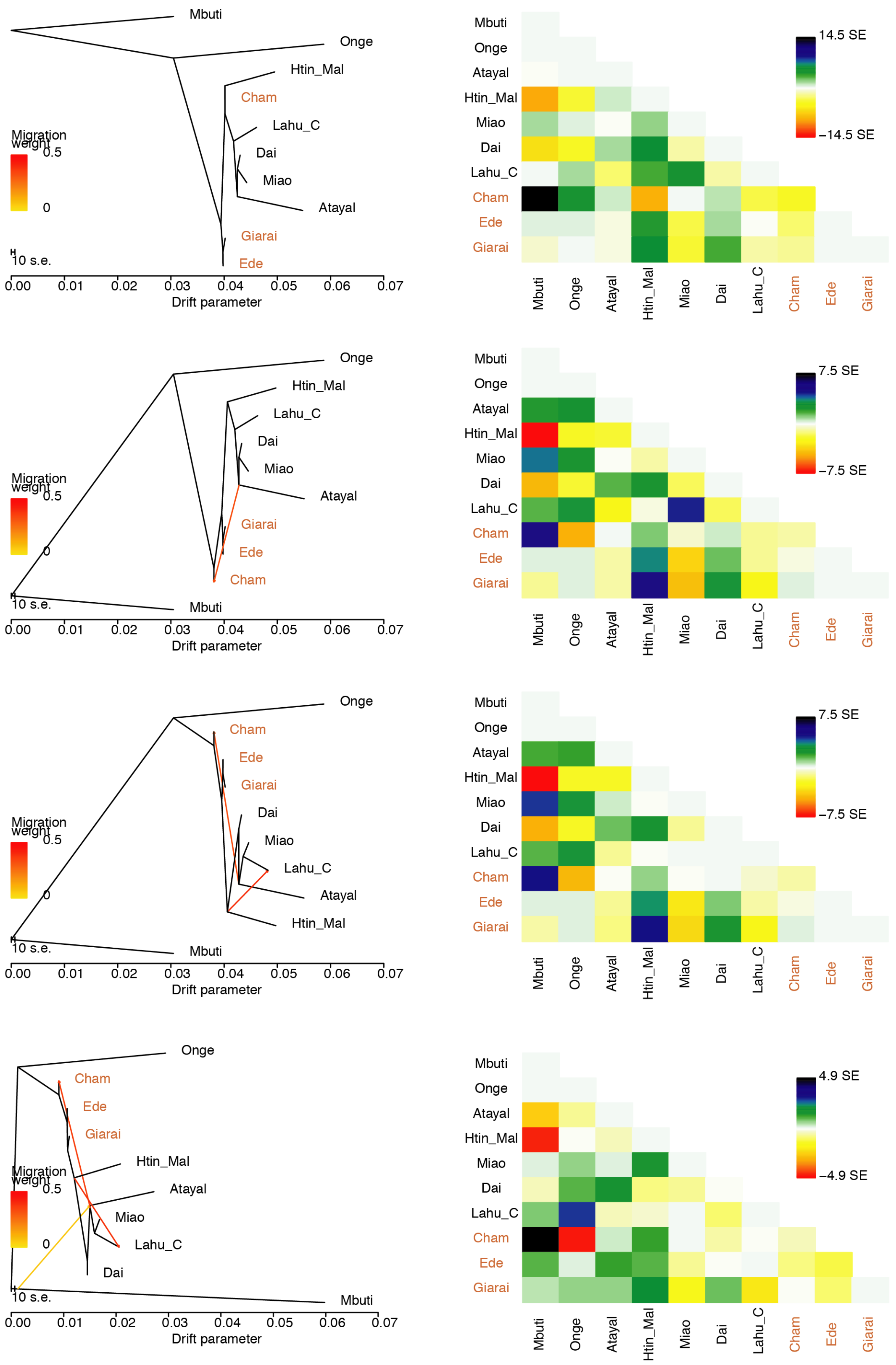


**Fig. S18. TreeMix results for the Vietnamese AN groups with 0 to 3 migrations.**

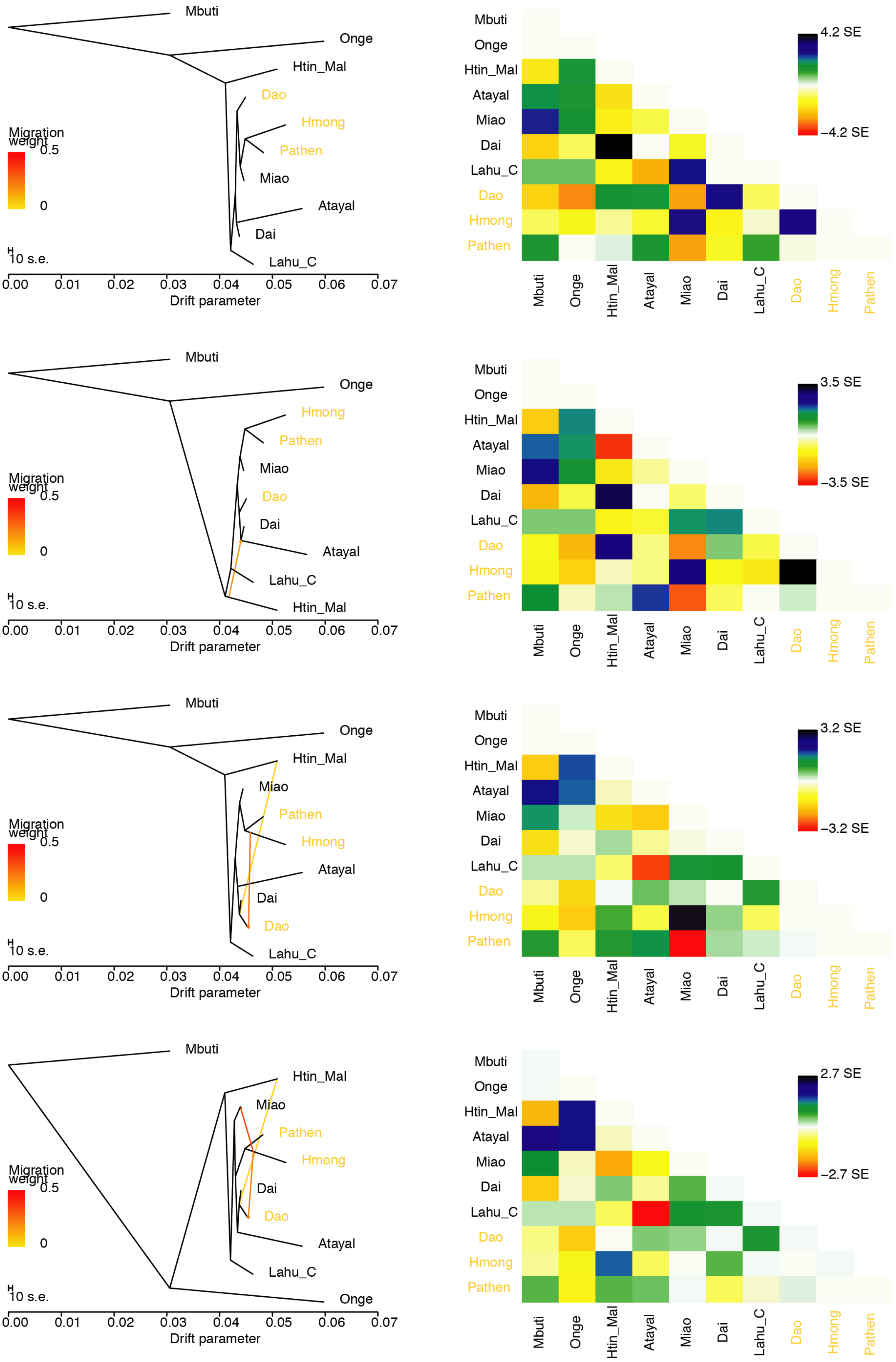


**Fig. S19. TreeMix results for the Vietnamese HM groups with 0 to 3 migrations.**

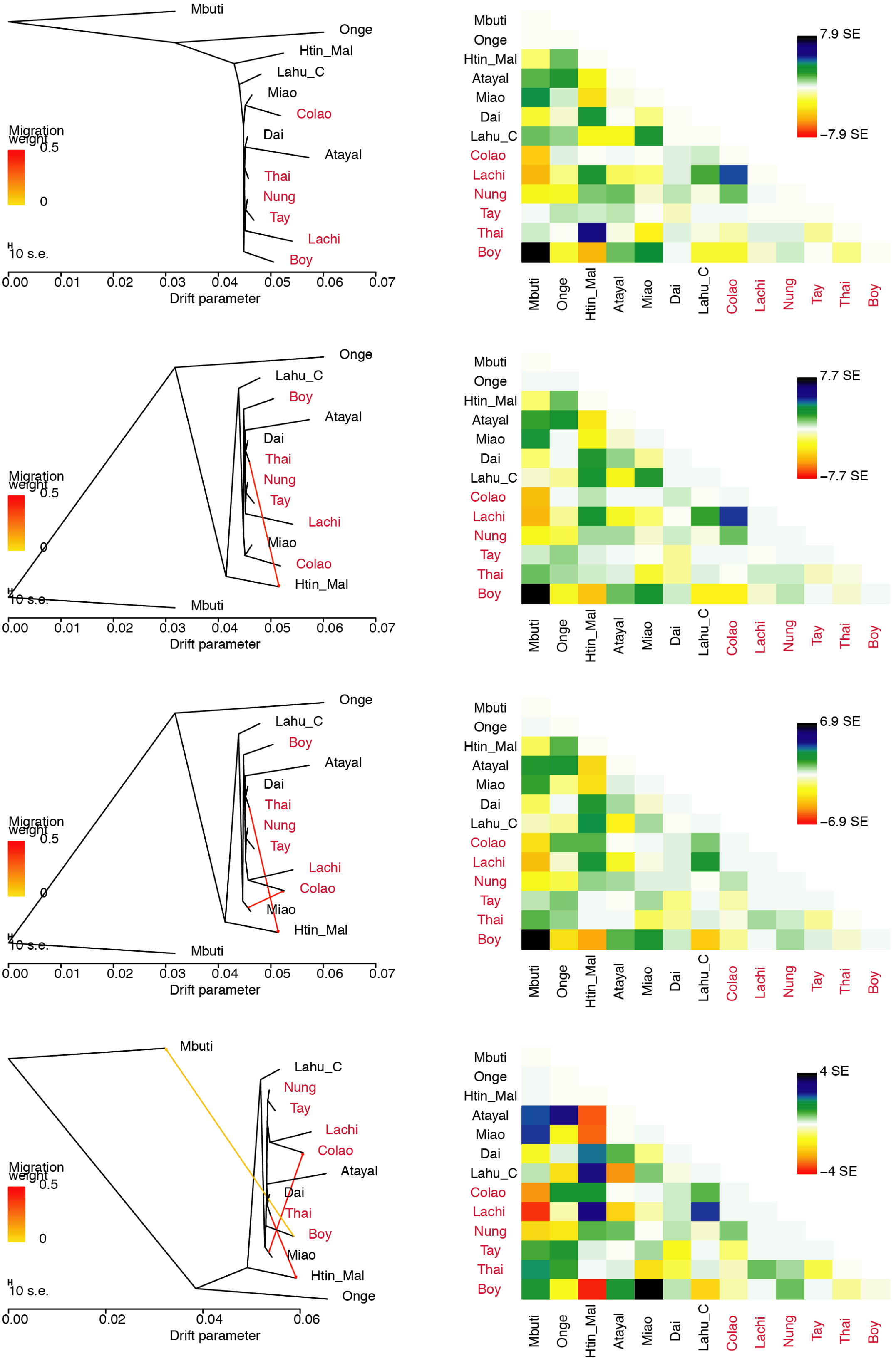


**Fig. S20. TreeMix results for the Vietnamese TK groups with 0 to 3 migrations.**

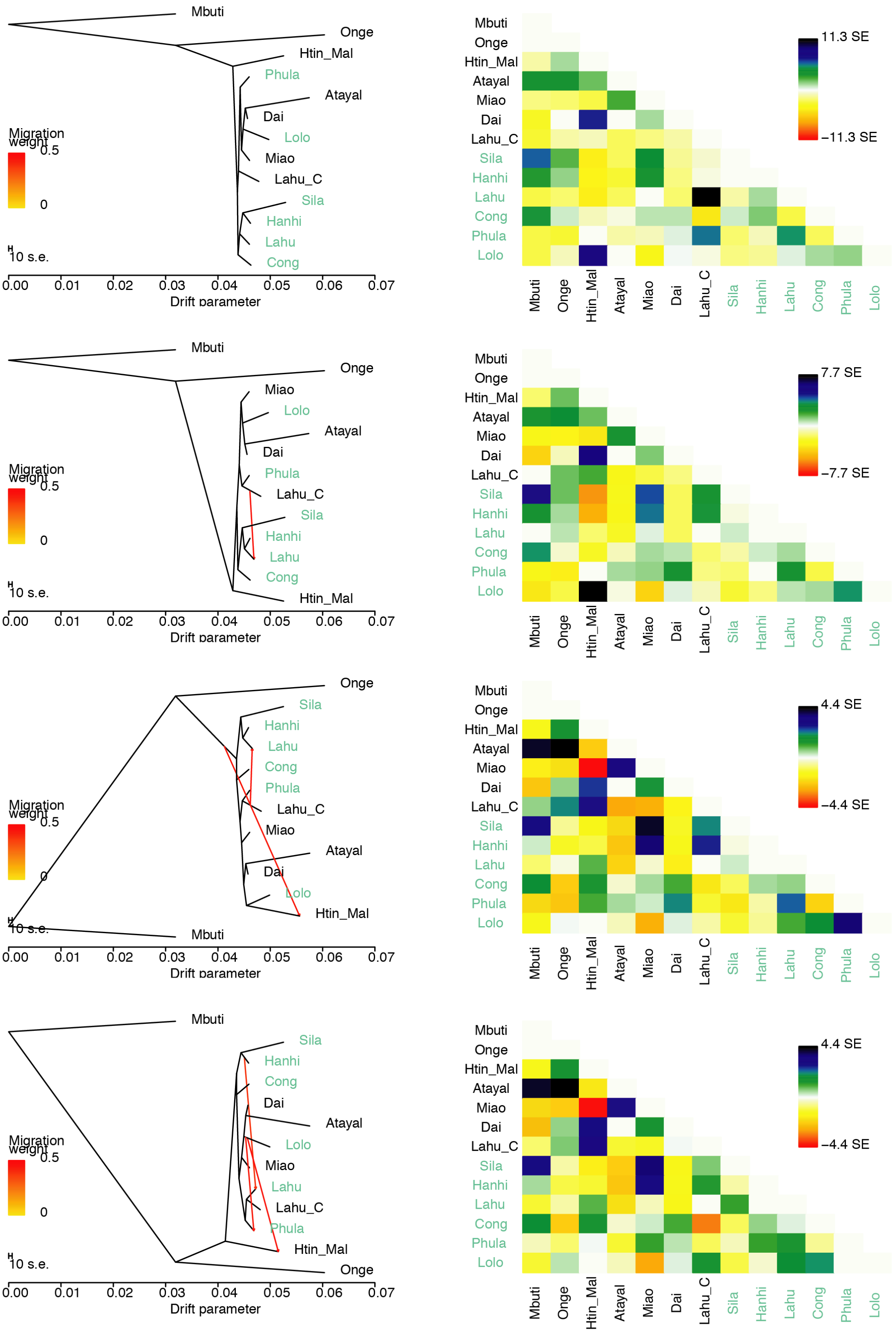


**Fig. S21. TreeMix results for the Vietnamese ST groups with 0 to 3 migrations.**

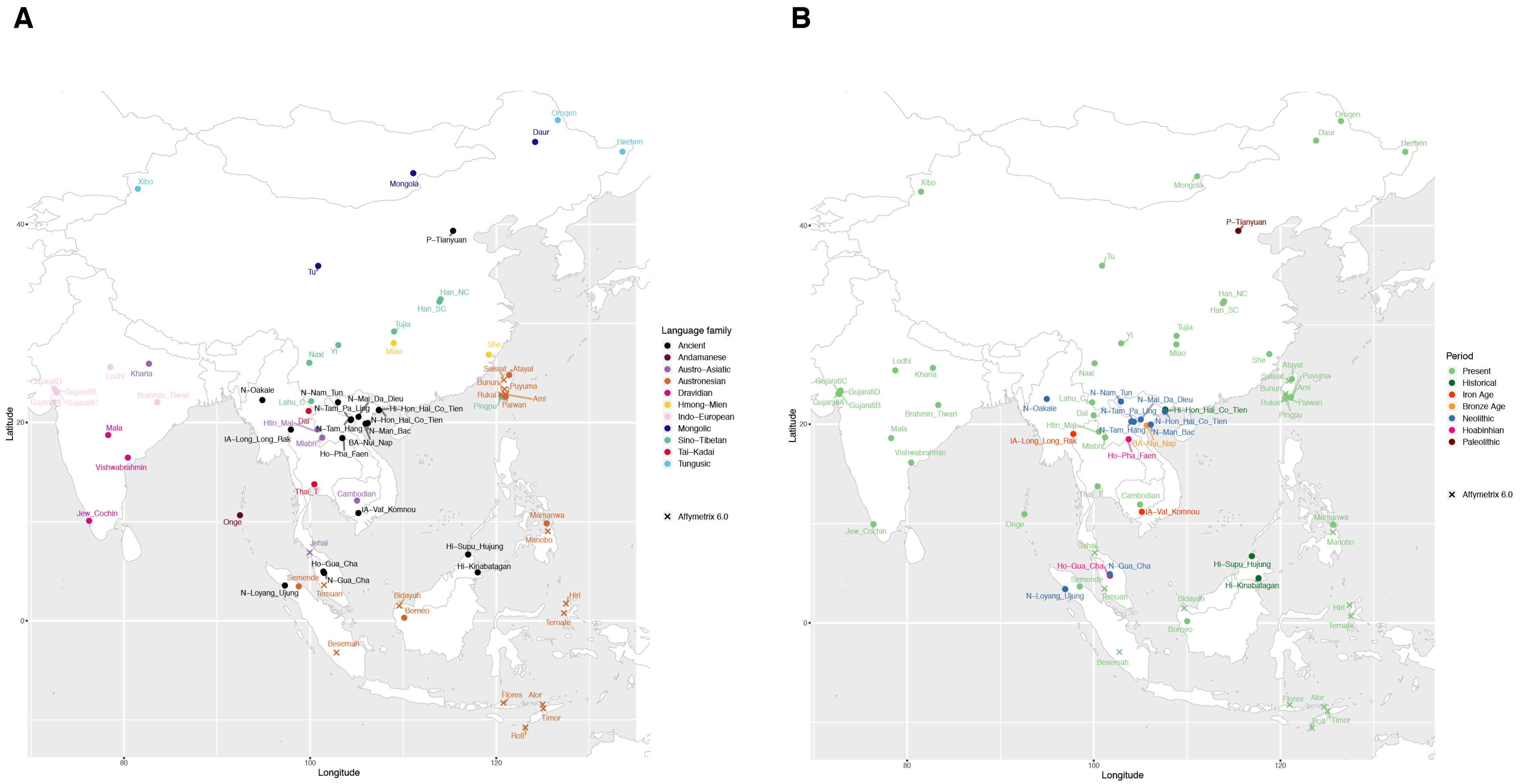


**Fig. S22. Sample information map.**
